# *PDF2.2* links constitutive defensin expression to disease resistance in *Arabidopsis*

**DOI:** 10.64898/2026.10.04.756585

**Authors:** Shailja Singh

## Abstract

**Background:** Plant defensins are small cysteine-rich antimicrobial peptides. In *Arabidopsis thaliana*, the inducible Class I gene *AtPDF1.2a* is widely used as a transcriptional marker of jasmonate/ethylene immunity, whereas the constitutively expressed Class II members have received less functional attention. AtPDF2.2 was compared with *AtPDF1.2a* during infection of *Arabidopsis* by the vascular fungal pathogen *Verticillium longisporum*, the necrotrophic fungus *Sclerotinia sclerotiorum* and the bacterium *Pseudomonas syringae* pv. tomato DC3000.

**Results:** Profiling of the AtPDF family showed that *AtPDF2.2*, together with the Class II genes *AtPDF2.3* and *AtPDF2.5*, was significantly reduced during *V. longisporum* infection, whereas *AtPDF1.2a* was induced. A promoter-reporter line showed a corresponding decrease in *AtPDF2.2* promoter activity during *V. longisporum* infection, strongest in infected roots. KD-*pdf2.2* plants contained more *V. longisporum* DNA, whereas OE-PDF2.2 plants contained less. The same reciprocal disease pattern was observed during *S. sclerotiorum* and Pst DC3000 infection. Histochemical Prom::*AtPDF2.2*::*GUS* reporter signal decreased during *S. sclerotiorum* infection but increased following Pst DC3000 infection. Basal *AtPDF2.2* expression was lower in JA- and ET-signaling mutants and higher in the SA-deficient *sid2* background, while the analyzed promoter lacked the GCC-box that mediates inducible *AtPDF1.2a* regulation. *AtPDF1.2a* overexpression reduced symptoms across the three pathosystems, but loss of *AtPDF1.2a* increased disease only during *V. longisporum* infection under the conditions tested and did not alter Pst DC3000 DNA abundance. *AtPDF2.2* overexpression was not associated with reduced vegetative growth under the conditions tested, whereas *AtPDF1.2a* overexpression reduced rosette size and leaf number.

**Conclusions:** *AtPDF2.2* is a constitutively expressed Class II defensin whose expression level is associated with disease outcome across two fungal pathogens and a bacterium. Its promoter shows tissue-dependent regulation during *V. longisporum* infection, while histochemical reporter signal decreases during *S. sclerotiorum* infection and increases during Pst DC3000 infection. This regulatory behavior differs from that of the inducible reference gene *AtPDF1.2a*. The reciprocal disease phenotypes of the *AtPDF2.2* transgenic backgrounds support a protective association that may involve direct antimicrobial activity, modulation of host responses or both. The absence of reduced vegetative growth in OE-PDF2.2 under the conditions tested supports further evaluation of *PDF2*-type genes in *Brassica*.

## Background

Plant defensins are small cysteine-rich antimicrobial peptides with roles in plant immunity as well as in development, reproduction and metal tolerance [1]. They share a conserved cysteine-stabilized alpha-beta fold but differ substantially in sequence, expression pattern and biological activity. The *Arabidopsis thaliana* family comprises fifteen loci assigned to three structural classes on the basis of their predicted mature peptides [2], whereas the allotetraploid *Brassica napus* genome contains 37 PDF genes [3]. This diversity extends to their transcriptional regulation. Class I genes such as *AtPDF1.2a* are expressed at low basal levels and are strongly induced through jasmonic acid (JA) and ethylene (ET) signaling during pathogen challenge [4–6], whereas several Class II members are expressed constitutively.

The distinction between inducible and constitutive expression may reflect different modes of antimicrobial protection. In radish, pre-existing defensins accumulate in peripheral seed tissues and are released during germination at concentrations sufficient to inhibit fungal growth [7]. Antifungal activity has also been demonstrated for PDF2-type proteins: recombinant *Arabidopsis AtPDF2.3* inhibits *Verticillium dahliae* and other phytopathogenic fungi *in vitro* [8], and *MaPDF2.2* from banana inhibits *Fusarium oxysporum* f. sp. cubense [9]. Constitutively expressed defensins could therefore contribute to protection without requiring strong transcriptional induction. Their functional importance in intact plants, however, remains less well characterized than that of the inducible PDF1.2 group.

The contribution of constitutively expressed defensins to resistance is particularly relevant during infections that extensively alter host transcription. *Verticillium longisporum* is a hybrid vascular fungus adapted to *Brassica*ceae and an important pathogen of oilseed rape (*B. napus*) [10, 11]. Following root infection, the fungus enters the vascular system and spreads systemically. Colonization is accompanied by changes in the apoplastic proteome, cell-wall properties, hormone signaling and defense-associated transcription [12–14]. Host susceptibility can also depend on plant genes that facilitate infection or disease development [15]. In the *V. longisporum* interaction, loss of *CRT1a* or *BnHva22c* reduces susceptibility [16, 17]. Whether constitutively expressed antimicrobial peptides are themselves altered during this interaction, and how such changes relate to disease outcome, has not been established.

This study examined the relationship between defensin transcription and disease resistance by comparing the constitutively expressed Class II gene *AtPDF2.2* with the inducible Class I reference *AtPDF1.2a*. Analyses of transcriptional responses, *AtPDF2.2* promoter regulation and disease-associated phenotypes during *V. longisporum* infection were combined with comparisons of the basal regulatory contexts of the two genes. Functional and promoter-reporter comparisons during *Sclerotinia sclerotiorum* and *Pseudomonas syringae* pv. tomato DC3000 infection assessed whether the relationship between expression behavior and disease outcome was maintained across pathogens with different infection strategies.

## Results

### Transcriptional reprogramming of the *AtPDF* family during *Verticillium longisporum* infection

The *Arabidopsis thaliana* plant defensin (*AtPDF*) family comprises fifteen members in three structural classes defined by their predicted mature peptides (Fig. 1A): seven Class I loci (*AtPDF1.1*, the near-identical *AtPDF1.2a/1.2b/1.2c*, *AtPDF1.3*, *AtPDF1.4* and *AtPDF1.5*), six Class II loci (*AtPDF2.1-2.6*) and two Class III genes (*AT5G38330* and *AT4G30070*) [2]. The expression analysis comprises thirteen profiles, including *AtPDF1.2a* but not separate profiles for *AtPDF1.2b* and *AtPDF1.2c, which* encode essentially identical mature peptides.

**Figure 1.**
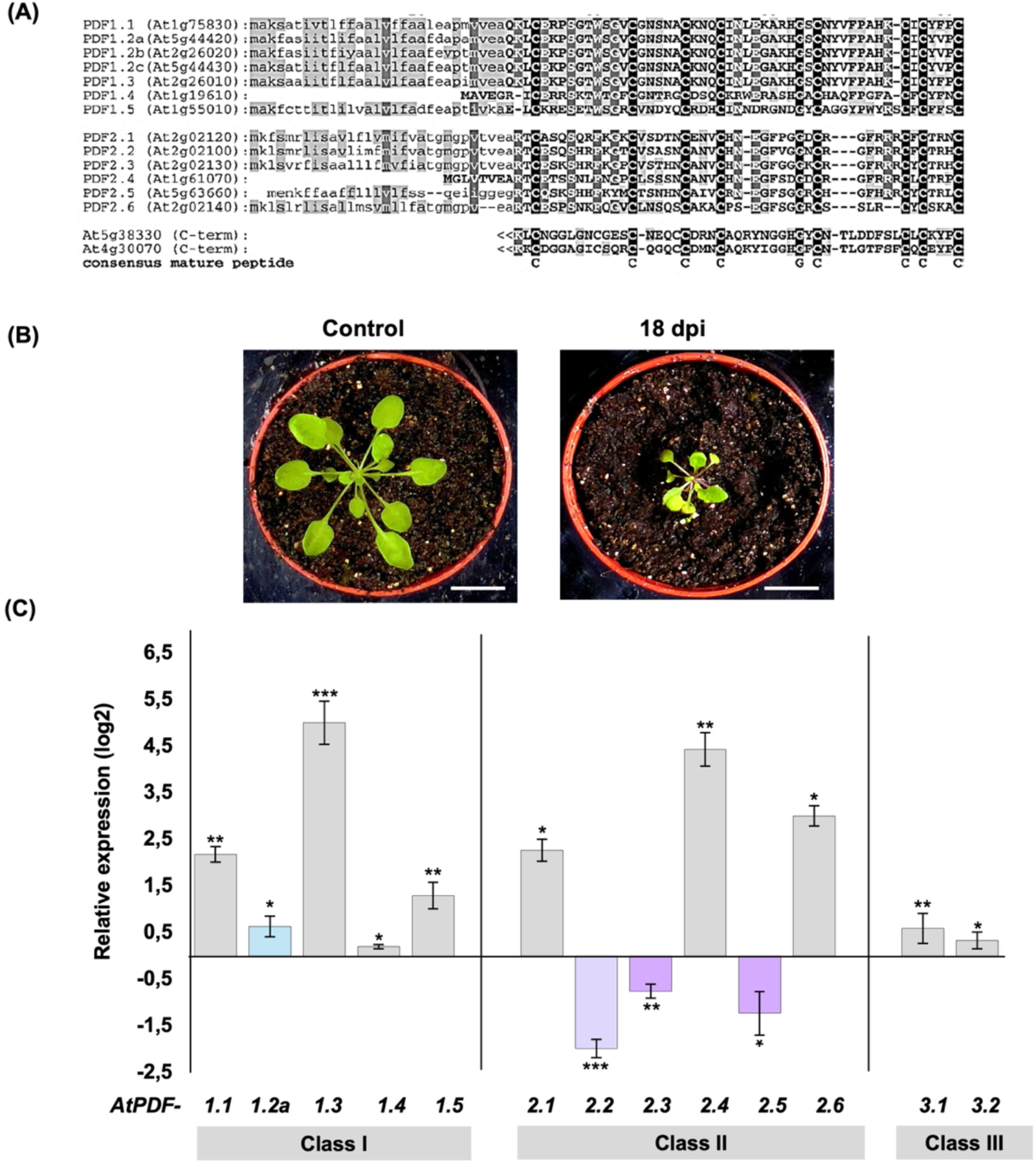
Expression of the *AtPDF* gene family during *Verticillium longisporum* infection. (A) Alignment of predicted mature peptides of the *Arabidopsis* plant defensin (*AtPDF*) family: Class I (*AtPDF1.1-1.5*), Class II (*AtPDF2.1-2.6*) and Class III (*AT5G38330*, *AT4G30070*; C-terminal domains shown), with the conserved cysteine consensus below. (B) Soil-grown Col-0 rosettes, mock-treated and Vl43-inoculated, at 18 days post-inoculation (dpi). Bars = 1 cm. (C) Transcript levels of *AtPDF* family members in Vl43-infected *in vitro* roots at 6 dpi, expressed as log₂ relative expression normalized to ACT2. Class III members are labeled 3.1 (*AT5G38330*) and 3.2 (*AT4G30070*). Thirteen transcript profiles are shown, including *AtPDF1.2a* but not separate profiles for *AtPDF1.2b* and *AtPDF1.2c*. Transcript levels were determined by RT-qPCR relative to *ACT2* (AT3G18780). Data are mean ± SD of three to four biological replicates. Asterisks indicate significant differences from the mock control (Student’s t-test): *\*P < 0.05, **P < 0.01, ***P < 0.005*.

Soil-grown Col-0 plants infected with *V. longisporum* isolate Vl43 were visibly stunted by 18 days post-inoculation (dpi) relative to mock-treated plants (Fig. 1B), consistent with the established *Arabidopsis*-Vl43 pathosystem [18, 19]. Expression profiling in infected roots at 6 dpi revealed contrasting responses within the defensin family. *AtPDF1.2a* was induced from a low basal level, whereas *AtPDF2.2*, together with *AtPDF2.3* and *AtPDF2.5*, was significantly reduced (Fig. 1C).

*AtPDF1.2a* induction is consistent with its established JA/ET-responsive expression [4–6]. The simultaneous reduction of several Class II genes identified a distinct transcriptional response within the same defensin family that included *AtPDF2.2*.

### *AtPDF2.2* promoter activity is dynamically reprogrammed during *Verticillium longisporum* colonization

Prom::*AtPDF2.2*::*GUS* plants showed reporter activity throughout untreated seedlings, including strong staining in the root, consistent with the constitutive expression of *AtPDF2.2* (Fig. 2A, B). Following *V. longisporum* infection, however, reporter staining was markedly reduced in roots at 6 dpi (Fig. 2B). A transcriptional component was supported by the time course: endogenous *AtPDF2.2* and *GUS* reporter transcripts followed closely similar trajectories between 3 and 9 dpi (Fig. 2C).

**Figure 2.**
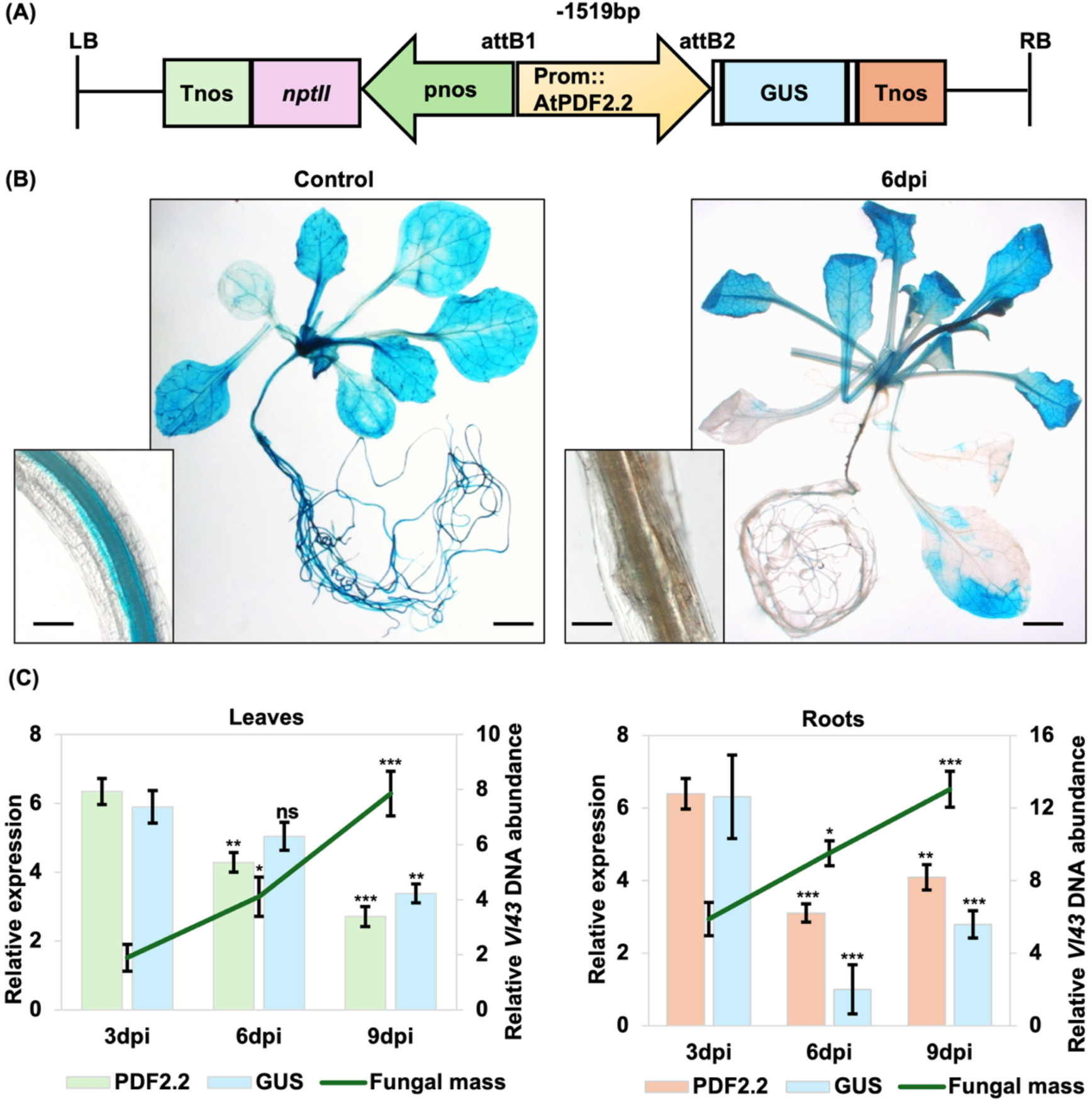
*AtPDF2.2* promoter activity and fungal biomass during *Verticillium longisporum* infection. (A) Schematic of the Prom::*AtPDF2.2*::*GUS* reporter T-DNA in pGWB433. A 1,519-bp fragment upstream of the *AtPDF2.2* start codon was introduced by Gateway LR recombination between attB1 and attB2, upstream of the promoterless β-glucuronidase (*GUS*) reporter, followed by a nopaline synthase (NOS) terminator. A pnos-*nptII*-Tnos cassette confers kanamycin selection. (B) *GUS* staining of Prom::*AtPDF2.2*::*GUS* plants, control and 6 dpi with *V. longisporum*, with root close-ups (insets). Bars = 2 mm (seedling), 50 µm (insets). (C) Transcript levels of *AtPDF2.2* and *GUS* (left axis) and relative Vl43 DNA abundance as a measure of fungal biomass (right axis, green line) in leaves and roots at 3, 6 and 9 dpi. Transcript levels were determined by RT-qPCR relative to *ACT2*, with each series quantified independently. Data are mean ± SD of three independent lines, each line a mean of three to four biological replicates. Asterisks indicate significant differences from the 3 dpi value of the same series (Student’s t-test): *\*P < 0.05, **P < 0.01, ***P < 0.005; ns, not significant*.

The response was strongly tissue- and time-dependent. In roots, both endogenous *AtPDF2.2* and reporter transcripts declined to their lowest measured levels at 6 dpi and subsequently increased by 9 dpi. In leaves, by contrast, both transcripts continued to decline through 9 dpi. *V. longisporum* enters through the root and elicits pronounced early transcriptional and hormone-associated responses in below-ground tissues, while systemic responses develop as vascular colonisation progresses [13, 14, 20]. Root-specific transcriptional reprogramming during *V. longisporum* infection has likewise been reported for multiple immunity-associated pathways [20].

The transcriptional pattern also differed from pathogen accumulation. Relative Vl43 DNA abundance increased between 3 and 9 dpi in both roots and leaves, whereas *AtPDF2.2* promoter output reached a minimum and began to recover in roots during the same interval (Fig. 2C). The reduction in *AtPDF2.2* expression therefore did not simply track increasing fungal biomass. Instead, the close correspondence between endogenous *AtPDF2.2* and Prom::*AtPDF2.2*::*GUS* indicates that regulatory information contained within the analysed 1,519-bp upstream region contributes to the infection-associated response. Together, these data identify the *AtPDF2.2* promoter as a dynamically regulated component of the *V. longisporum* interaction, with distinct temporal behaviour in roots and leaves.

### *AtPDF1.2a* and *AtPDF2.2* transgenic lines show reciprocal *Verticillium longisporum* phenotypes

Disease responses were compared between *AtPDF1.2a* knock-out (KO-*pdf1.2a*) and overexpression (OE-PDF1.2a) lines and *AtPDF2.2* knock-down (KD-*pdf2.2*) and overexpression (OE-PDF2.2) lines (Fig. 3). During infection, *AtPDF1.2a* was induced whereas *AtPDF2.2* was reduced in wild-type plants, consistent with the family-wide expression analysis (Fig. 3C, D).

**Figure 3.**
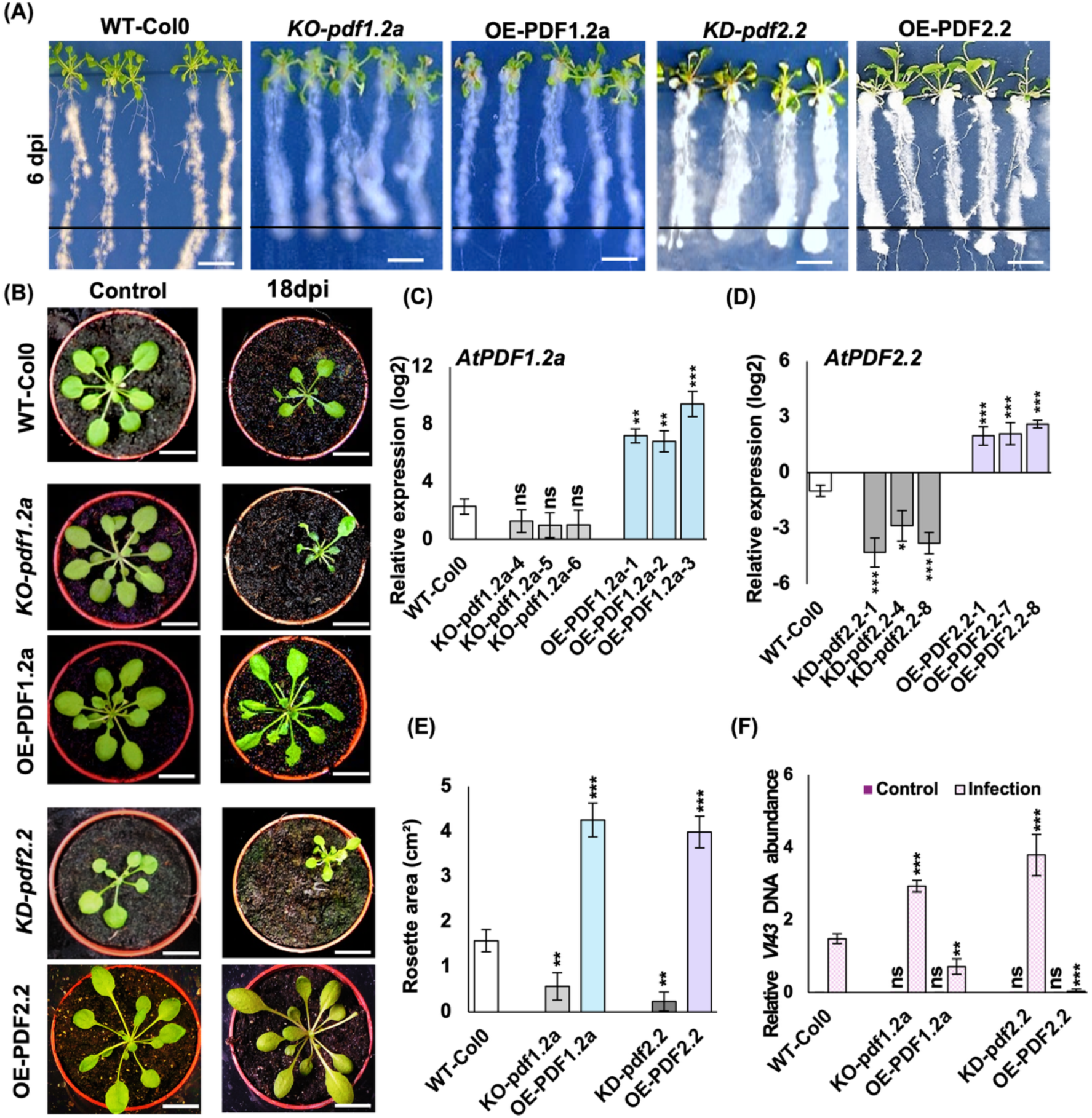
Phenotypes and transcript levels of *AtPDF1.2a* and *AtPDF2.2* transgenic lines under *Verticillium longisporum* infection. (A) Root colonization at 6 dpi of *in vitro*-grown wild-type (WT-Col-0), KO-*pdf1.2a*, OE-PDF1.2a, KD-*pdf2.2* and OE-PDF2.2 plants. The horizontal line marks the inoculation front. Bars = 1 cm. (B) The same genotypes grown in soil under control and Vl43-infected conditions at 18 dpi. Bars = 1 cm. (C, D) Transcript levels of *AtPDF1.2a* (C) and *AtPDF2.2* (D) during Vl43 infection, determined by RT-qPCR, normalized to *ACT2* and expressed as log₂ relative expression, with the three lines per genotype and infected WT-Col-0 plotted individually. T-DNA insertions were confirmed by genotyping (Table S3). Data are mean ± SD of three to four biological replicates per line. (E) Rosette area (cm²) of infected plants at 18 dpi. These values are not directly comparable with the developmental rosette areas of non-infected plants in Fig. S1E. (F) Relative Vl43 DNA abundance in control and infected plants. (E, F) Data are mean ± SD across three lines (n = 3); each line value represents the mean of the biological replicates within that line. Asterisks indicate significant differences from the correspondingly treated WT-Col-0 (Student’s t-test): *\*P < 0.05, **P < 0.01, ***P < 0.005; ns, not significant*.

Root colonization at 6 dpi and disease symptoms at 18 dpi were greater in KO-*pdf1.2a* and KD-*pdf2.2* plants and reduced in both overexpression backgrounds relative to infected wild type (Fig. 3A, B). Infected KO-*pdf1.2a* and KD-*pdf2.2* plants also had smaller rosettes, whereas OE-PDF1.2a and OE-PDF2.2 plants retained larger rosettes (Fig. 3E). Relative Vl43 DNA abundance followed the same pattern, increasing in KO-*pdf1.2a* and KD-*pdf2.2* plants and decreasing in the corresponding overexpression backgrounds (Fig. 3F).

Thus, opposite infection-induced transcriptional responses did not correspond to opposite disease phenotypes. For *AtPDF2.2*, transcript abundance decreased during infection, yet KD-*pdf2.2* plants contained more fungal DNA and OE-PDF2.2 plants contained less. The infection-associated decline in *AtPDF2.2* expression therefore occurred alongside a positive association between higher expression and resistance.

The transgenic lines also differed under non-infected conditions (Fig. S1). OE-PDF2.2 plants did not show reduced vegetative growth: primary roots and rosettes were larger than those of wild type at the measured stages, whereas bolting time and leaf number were unchanged. In contrast, OE-PDF1.2a plants had smaller rosettes and fewer leaves.

KD-*pdf2.2* plants also showed developmental changes, including earlier bolting, fewer leaves and reduced rosette area. High or constitutive expression of defense components can impose physiological costs, although these effects vary with the gene and expression system [21–23]. Because the genotypes differed developmentally before infection, rosette area was interpreted together with Vl43 DNA abundance, which provided a DNA-based estimate of fungal biomass. Under the conditions tested, increased resistance in OE-PDF2.2 was not accompanied by the reduction in vegetative growth observed in OE-PDF1.2a.

### *AtPDF2.2* has a basal regulatory context distinct from inducible *AtPDF1.2a*

Comparisons in hormone-signaling mutants revealed distinct basal regulatory responses of AtPDF2.2 and the inducible reference *AtPDF1.2a*. Basal *AtPDF2.2* expression was significantly lower in mutants affecting JA or ET signaling, with the strongest decrease in *coi1-16 ein2-1*, and was higher in the SA-deficient *sid2* mutant (Fig. 4). In contrast, basal *AtPDF1.2a* expression was largely unchanged in the JA- and ET-related mutants and increased in *sid2*. These mutant responses are consistent with positive JA/ET-associated and negative SA-associated influences on basal *AtPDF2.2* expression. Interactions among SA, JA and ET are strongly context dependent in plant immunity [24–26]. Together, these data place *AtPDF2.2* in a basal regulatory context influenced by JA-, ET- and SA-associated pathways and provide a framework for examining the additional regulation associated with infection.

**Figure 4.**
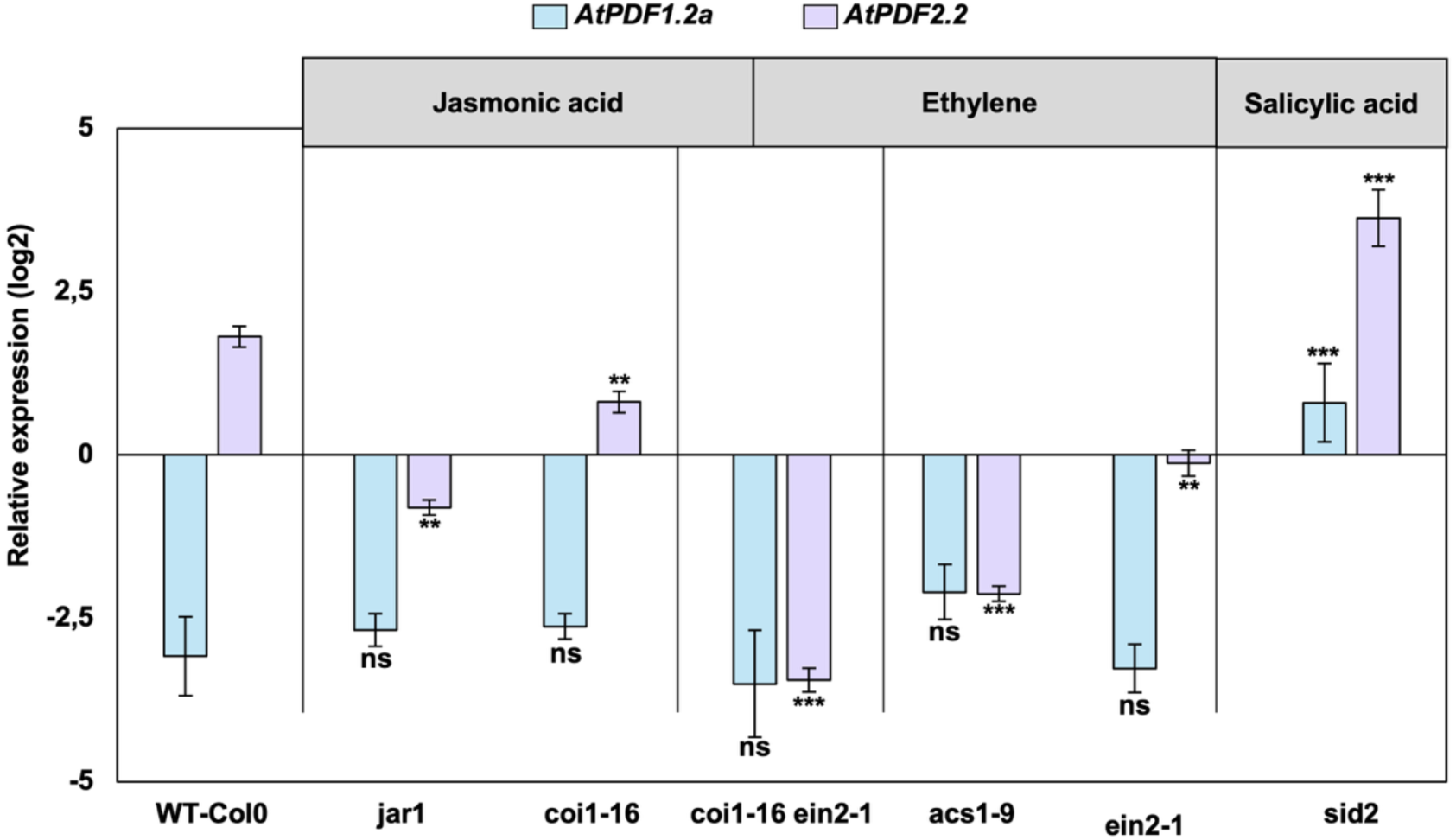
Basal *AtPDF1.2a* and *AtPDF2.2* expression in hormone-signaling mutants. Transcript levels of *AtPDF1.2a* and *AtPDF2.2* in non-infected leaves of Col-0 and mutants affected in jasmonic acid signaling (*jar1*, *coi1-16*), combined jasmonic acid and ethylene signaling (*coi1-16 ein2-1*), ethylene biosynthesis and signaling (*acs1-9*, *ein2-1*), and salicylic acid biosynthesis (*sid2*). Transcript levels were determined by RT-qPCR and are expressed as log₂ relative expression normalized to *ACT2*. Data are mean ± SD of three to four biological replicates. Asterisks indicate significant differences from Col-0 (Student’s t-test): *\*P < 0.05, **P < 0.01, ***P < 0.005; ns, not significant*.

The transcriptional contexts of the two genes were also distinct. *AtPDF1.2a* co-expressed with PR proteins, chitinase, beta-1,3-glucanase and components of the ERF/ORA59 defense network, whereas *AtPDF2.2* was preferentially associated with circadian-and abiotic-stress-responsive genes, including *TOC1*, *ELF3*, *ELF4*, *GRP7*, *RCI2A* and *PIP2;5* (Fig. S2). GO enrichment similarly separated defense- and fungus-associated processes among *AtPDF1.2a* partners from temperature, circadian and abiotic-stress responses among *AtPDF2.2* partners (Fig. S3A).

The promoter architectures were likewise different. The *AtPDF1.2a* promoter contains the GCC-box through which ORA59 integrates JA and ET signals during defense induction [27, 28], whereas the analyzed *AtPDF2.2* promoter lacks this element and contains low-temperature- and circadian-associated motifs (Fig. S3B). These features distinguish the promoter contexts of the two genes and provide candidate regulatory elements for future functional analysis of *AtPDF2.2* regulation.

Altered *AtPDF2.2* expression was accompanied by broader changes in hormone-associated transcripts. JA-biosynthesis genes (*LOX3*, *AOC3*, *OPR3*), ET-signaling genes (*ETR2*, *EIN3*) and SA-associated genes (*PR1*, *WRKY70*, *WRKY53*) generally changed in opposite directions in KD-*pdf2.2* and OE-PDF2.2 during *V. longisporum* infection (Fig. S4B). Changes in KO-*pdf1.2a* and OE-PDF1.2a were less consistently related to gene expression level (Fig. S4A). These patterns place the *AtPDF2.2* transgenic backgrounds within broader shifts in hormone-associated transcription that accompany the altered infection phenotypes.

Several Class II genes also changed in the *AtPDF2.2* transgenic lines. By 12 dpi with *V. longisporum*, *AtPDF2.1*, *AtPDF2.2*, *AtPDF2.3* and *AtPDF2.5* transcripts were lower in KD-*pdf2.2* and higher in OE-PDF2.2 than in infected wild type (Fig. S5). Earlier responses varied with the gene and sampling time. Manipulation of *AtPDF2.2* was therefore accompanied by coordinated changes in related Class II genes. This broader response is compatible with shared regulation within the Class II group as well as effects linked to the transgenic background or differences in infection progression. Functional diversification nevertheless exists within this group: *AtPDF2.1* has been linked to ammonium metabolism and to promoter activity in nematode-induced syncytia [29, 30]. Together, Fig. 4 and Figs. S2-S5 place *AtPDF2.2* in a regulatory context distinct from *AtPDF1.2a*.

### *AtPDF2.2* transgenic lines show reciprocal responses to *Sclerotinia sclerotiorum* as reporter signal declines

During infection with the necrotrophic fungus *S. sclerotiorum*, KD-*pdf2.2* plants showed greater whole-plant disease severity, whereas OE-PDF2.2 plants were less affected than wild type (Fig. 5A, C). Detached leaves showed the same relationship, with larger lesions in KD-*pdf2.2* and smaller lesions in OE-PDF2.2 at 24 hpi (Fig. 5B, D). Histochemical Prom::*AtPDF2.2*::*GUS* staining became progressively weaker between 6 and 24 hpi (Fig. 5E). Thus, a reduced reporter signal accompanied infection by a second fungus; in *S. sclerotiorum* this histochemical response provides a qualitative counterpart to the transcript-resolved *V. longisporum* time course.

**Figure 5.**
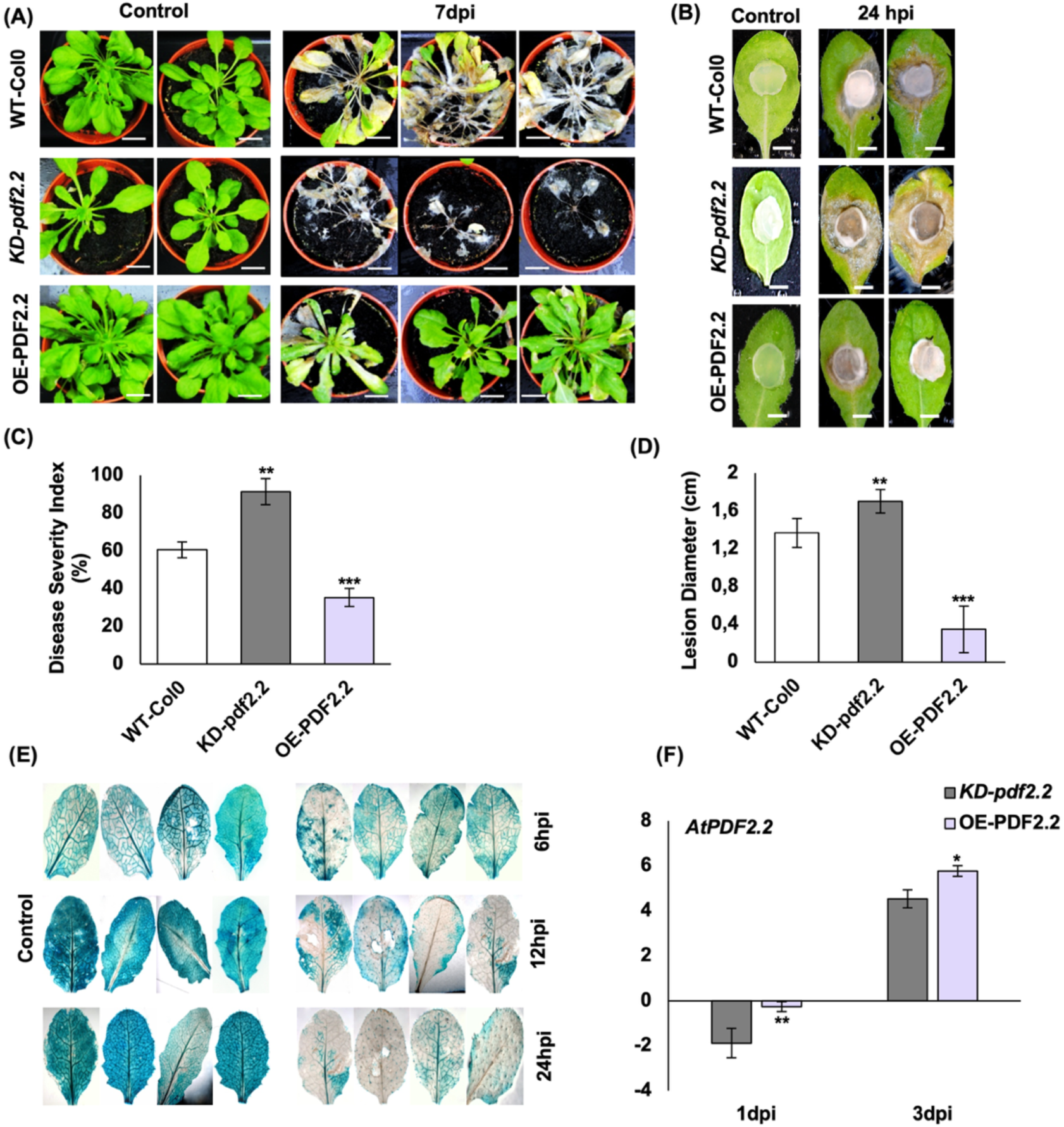
*AtPDF2.2* transgenic lines under *Sclerotinia sclerotiorum* infection. (A) Whole-plant phenotype of WT-Col-0, KD-*pdf2.2* and OE-PDF2.2 at 7 dpi and in the control. Bars = 1 cm. (B) Detached-leaf assay of the same genotypes at 24 h post-inoculation (hpi) and in the control. Bars = 1 cm. (C) Disease Severity Index (%) of whole plants scored at 7 dpi on the 0-6 scale defined in Methods. (D) Lesion diameter (cm) in the detached-leaf assay at 24 hpi. (C, D) Data are mean ± SD across three lines (n = 3); each line value represents the mean of the biological replicates within that line. (E) Prom::*AtPDF2.2*::*GUS* staining of control and infected leaves at 6, 12 and 24 hpi. Bars = 2 mm. (F) Transcript levels of *AtPDF2.2* in KD and OE lines at 1 and 3 dpi, determined by RT-qPCR, normalized to *ACT2* and expressed as log₂ relative expression. Data are mean ± SD of three to four biological replicates. Fungal biomass was not quantified in this pathosystem. (C, D, F) Asterisks indicate significant differences from the correspondingly treated WT-Col-0 (Student’s t-test): *\*P < 0.05, **P < 0.01, ***P < 0.005; ns, not significant*.

The corresponding *AtPDF1.2a* lines showed a different loss-of-function response. OE-PDF1.2a plants developed less severe whole-plant disease and smaller detached-leaf lesions, whereas KO-*pdf1.2a* did not differ significantly from wild type (Fig. S6). Increased expression of either defensin was therefore associated with reduced disease, but loss of *AtPDF1.2a* did not measurably alter *S. sclerotiorum* disease under the conditions tested.

### *AtPDF2.2* transgenic lines also show reciprocal responses to Pst DC3000 while reporter signal increases

Pst DC3000 produced stronger Prom::*AtPDF2.2*::*GUS* staining than the mock treatment, opposite to the reduced reporter signal observed during the fungal interactions (Fig. 6C). The disease phenotypes of the *AtPDF2.2* transgenic lines nevertheless followed the same direction as in the fungal interactions. KD-*pdf2.2* plants developed more severe symptoms and contained significantly more bacterial DNA, whereas OE-PDF2.2 plants showed reduced symptoms and lower bacterial DNA abundance (Fig. 6A, B, E).

**Figure 6.**
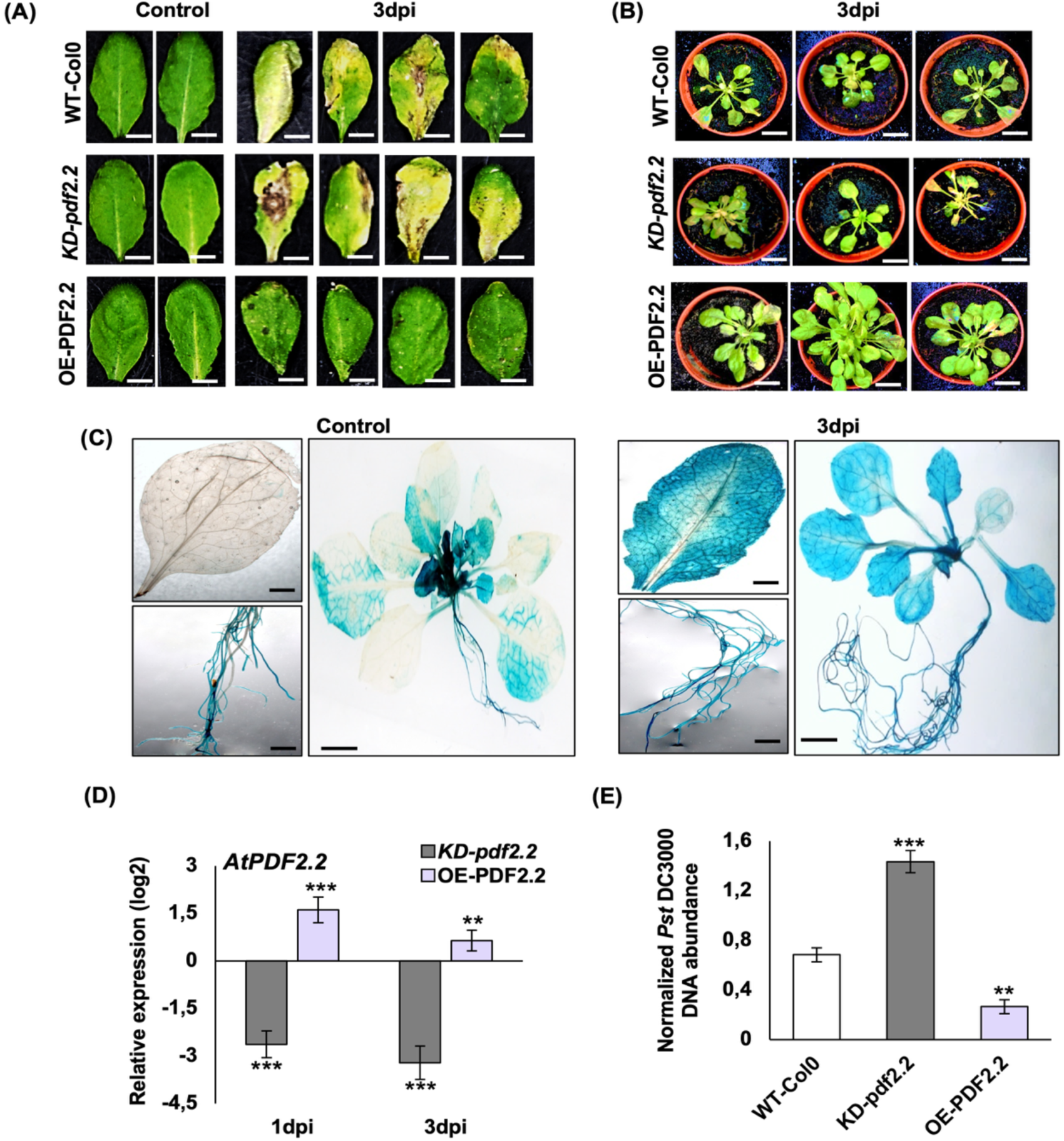
*AtPDF2.2* transgenic lines under *Pseudomonas syringae* pv. tomato DC3000 infection. (A) Detached-leaf symptoms of WT-Col-0, KD-*pdf2.2* and OE-PDF2.2 under control conditions and at 3 dpi. Bars = 1 cm. (B) Whole-rosette symptoms of the same genotypes under control conditions and at 3 dpi. Bars = 1 cm. (C) Prom::*AtPDF2.2*::*GUS* staining, control and 3 dpi. (D) Transcript levels of *AtPDF2.2* in KD and OE lines at 1 and 3 dpi, determined by RT-qPCR, normalized to *ACT2* and expressed as log₂ relative expression. Data are mean ± SD of three to four biological replicates. (E) Normalized Pst DC3000 DNA abundance. Data are mean ± SD across three lines (n = 3); each line value represents the mean of the biological replicates within that line. (D, E) Asterisks indicate significant differences from the correspondingly treated WT-Col-0 (Student’s t-test): *\*P < 0.05, **P < 0.01, ***P < 0.005; ns, not significant*.

OE-PDF1.2a plants also showed reduced symptoms following Pst DC3000 infection, whereas KO-*pdf1.2a* did not differ significantly from wild type (Fig. S7A, B). Bacterial DNA abundance was not significantly altered in either *AtPDF1.2a* genotype despite the differences in transcript abundance (Fig. S7C, D). In contrast, both symptoms and bacterial DNA abundance changed in the *AtPDF2.2* transgenic backgrounds.

Across the three pathosystems, KD-*pdf2.2* plants consistently showed greater disease, whereas loss of *AtPDF1.2a* produced a detectable phenotype only during *V. longisporum* infection under the conditions tested. The close relationship among *AtPDF1.2a*, *AtPDF1.2b* and *AtPDF1.2c*, which encode highly similar mature peptides, provides a plausible basis for the narrower loss-of-function phenotype across the other pathosystems. The diversity of the defensin family [2, 31] provides context for this possible redundancy, whose contribution may vary among pathosystems.

The three pathogens also differed in their effect on the *AtPDF2.2* reporter. The quantified *V. longisporum* time course and the histochemical *S. sclerotiorum* assay both showed a fungal-associated decrease, whereas Pst DC3000 produced stronger histochemical *GUS* staining. The contrasting reporter responses across pathogens indicate that promoter behavior is context dependent, with the fungal-associated decrease emerging as a specific feature rather than a general correlate of disease progression.

## Discussion

*AtPDF1.2a* has largely defined the transcriptional behavior associated with *Arabidopsis* defensins in immunity and served as an inducible reference for this canonical response. Its induction by pathogen challenge depends on JA/ET signaling and the ORA59-GCC-box module [4–6, 27, 28]. *AtPDF2.2* differs from this pattern. It is constitutively expressed, its basal transcript level is altered in JA-, ET-and SA-related mutant backgrounds, and its promoter activity decreases during *V. longisporum* infection. Nevertheless, KD-*pdf2.2* plants showed greater disease and OE-PDF2.2 plants showed reduced disease across all three pathosystems. Together, these findings separate infection-induced transcriptional behavior from the protective association of higher *AtPDF2.2* expression.

The promoter-reporter analysis further distinguishes *AtPDF2.2* from the inducible behavior of *AtPDF1.2a*. Similar temporal profiles of endogenous *AtPDF2.2* and Prom::*AtPDF2.2*::*GUS* transcripts indicate that the decrease during *V. longisporum* infection involves regulatory information contained within the analyzed 1,519-bp upstream region. The response was not uniform across tissues: promoter activity reached its lowest measured level earlier in roots and began to recover there while the decline continued in leaves and fungal abundance increased. Tissue-specific transcriptional and hormone responses have been reported during *V. longisporum* infection [13, 14, 20], providing context for this spatially resolved behavior. Histochemical reporter signal also decreased during *S. sclerotiorum* infection but increased after Pst DC3000 challenge. The *V. longisporum* time course provides quantitative resolution of this tissue-dependent behavior, while the additional pathosystems extend the comparison and show that the reporter response depends on pathogen context.

The fungal-associated promoter decrease therefore appears within a broader regulatory context. The hormone-mutant data define basal contributions of JA-, ET- and SA-associated pathways, while pathogen effectors and cross-kingdom small RNAs provide relevant precedents for infection-dependent modulation of host immune transcription [32, 33]. The correspondence between endogenous and reporter transcripts places at least part of the *V. longisporum*-responsive behavior within the analyzed promoter region and provides a basis for identifying the upstream signals involved.

Across the three pathosystems, the disease phenotype was more consistent than the direction of the reporter response: OE-PDF2.2 plants showed less disease and KD-*pdf2.2* plants more disease during *V. longisporum*, *S. sclerotiorum* and Pst DC3000 challenge. This convergence supports a broad protective association for higher *AtPDF2.2* expression across pathogens with distinct infection strategies. The different disease readouts, together with the absence of peptide-level measurements, leave room for both direct antimicrobial activity and modulation of host responses. The accompanying hormone-associated transcriptional changes further suggest that altered host signaling may form part of this protective phenotype.

Several related Class II genes changed in parallel with *AtPDF2.2* in KD-*pdf2.2* and OE-PDF2.2 plants (Fig. S5), indicating that *AtPDF2.2* manipulation occurs within a broader Class II transcriptional response. The KD phenotype is therefore interpreted in the context of this coordinated expression pattern, with reduced *AtPDF2.2* accompanied by reduced expression of related Class II genes. For *AtPDF1.2a*, the narrower loss-of-function phenotype across pathosystems is compatible with redundancy among the closely related PDF1.2 genes [2, 31]. Together, these observations support disease associations linked to defensin expression while emphasizing the contribution of the surrounding gene-family context.

Loss of *CRT1a* reduces susceptibility to *V. longisporum* and has been linked to enhanced ethylene-related transcription [16, 34]. In the *crt1a* knock-out, *AtPDF2.2* transcript was higher than in infected wild type, whereas *AtPDF1.2a* was unchanged (Fig. S8). This places *CRT1a* alongside the hormone-associated observations as a potential regulatory connection, while its relationship to disease and the basis of the expression change remain to be established. More broadly, loss of *CRT1a* or *BnHva22c* reduces susceptibility [16, 17], whereas the KD-*pdf2.2* background showed greater disease, placing the *AtPDF2.2*-associated phenotype in a different direction from these susceptibility-related genes. Other host determinants, including *ERECTA*-associated loci, also contribute to quantitative *V. longisporum* resistance [35], and the molecular basis of the *V. longisporum*-*Brassica*ceae interaction remains incompletely resolved [36].

The broad expression of PDF2-type genes in *B. napus* makes this crop a relevant system in which to examine whether the *Arabidopsis* phenotype is conserved. *B. napus* contains an expanded defensin repertoire, including peptides with demonstrated antimicrobial activity, and *BnaPDF2.2* homologues are expressed in several tissues [3, 37]. Defensin activity and regulation nevertheless differ among species: *TaPDF2.15* restricts *Phytophthora infestans* when transiently expressed [38], whereas the activity of *MtDef4.2* depends on its subcellular localization [39]. Evaluating whether the *AtPDF2.2* expression-disease relationship is conserved in *B. napus* would provide a direct test of its relevance beyond *Arabidopsis*. If conserved, regulated promoter strategies could then be considered as a way to maintain beneficial expression without relying on constitutive overexpression [40]. The relationships supported by the present study are summarized in Fig. 7.

**Figure 7.**
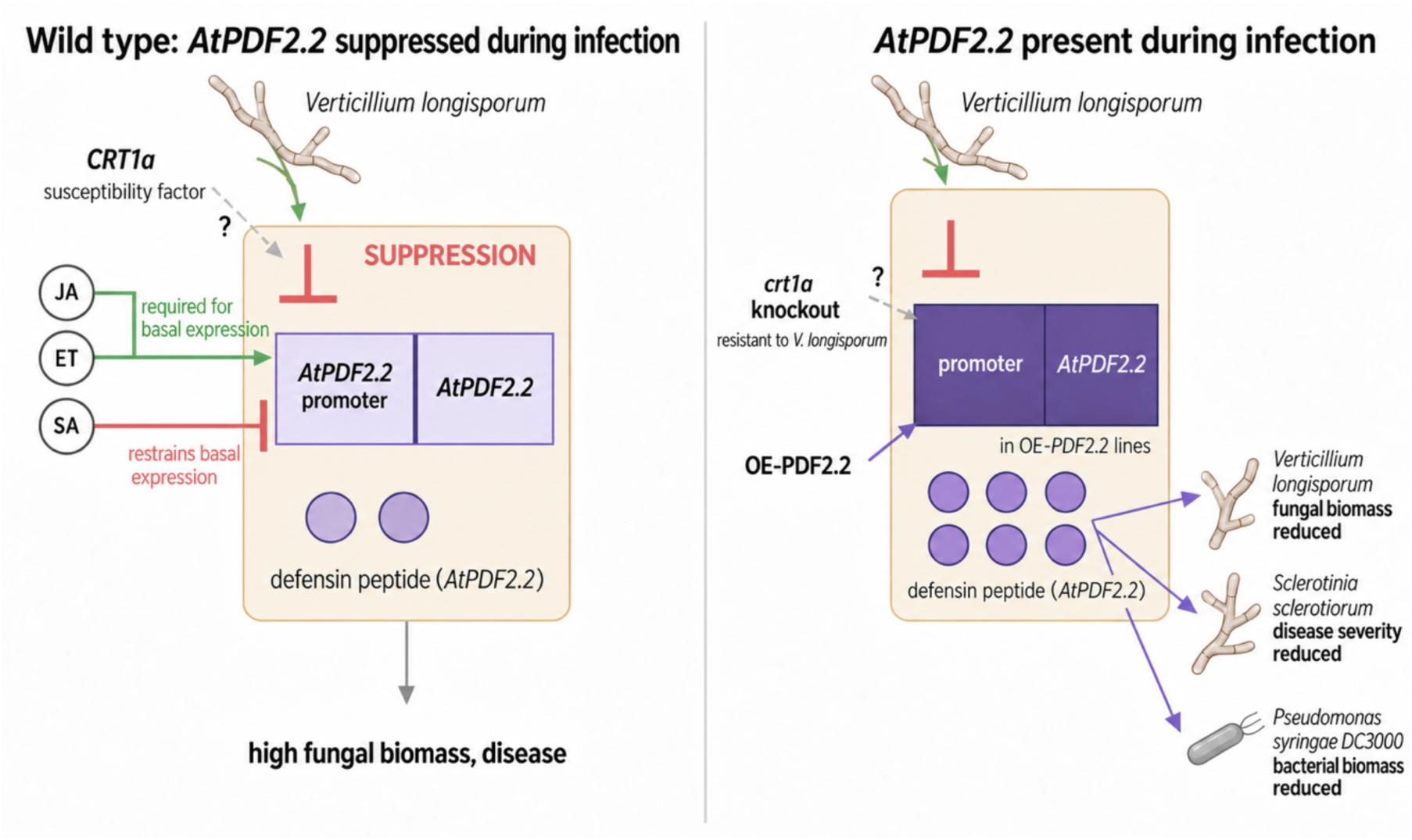
Working model of *AtPDF2.2* regulation and disease association. The left panel summarizes the regulatory relationships associated with basal and infection-responsive *AtPDF2.2* expression. Basal *AtPDF2.2* transcript abundance is lower in JA- and ET-signaling mutants and higher in the *sid2* background. The AtPDF2.2 promot*er-GUS reporter supports an* infection-associated decrease in promoter activity: endogenous *AtPDF2.2* and Prom::*AtPDF2.2*::*GUS* transcripts decrease with tissue-dependent kinetics during Verticillium longisporum infection. Histochemical GUS signal also decreases during *Sclerotinia sclerotiorum* infection but increases following *Pseudomonas syringae* pv. *tomato* DC3000 infection, indicating pathogen-dependent regulation of the *AtPDF2.2* promoter. The right panel summarizes disease associations in the transgenic backgrounds; overexpression was driven by the CaMV 35S promoter. Peptide symbols illustrate the proposed protective contribution rather than measured peptide abundance. KD-*pdf2.2* plants show greater disease or pathogen abundance, whereas OE-PDF2.2 plants show reduced disease or pathogen abundance across the three pathosystems. The KD phenotype is considered together with the accompanying changes in related Class II transcripts. Elevated *AtPDF2.2* transcript abundance in the *crt1a* background suggests a possible connection between *CRT1a* function and *AtPDF2.2* expression.

## Conclusions

*AtPDF2.2* is a constitutively expressed Class II defensin whose expression level is associated with disease outcome across *V. longisporum*, *S. sclerotiorum* and *Pst DC3000*. Its quantified promoter response during *V. longisporum* infection is tissue dependent, while histochemical reporter signal decreases during *S. sclerotiorum* infection and increases during bacterial infection. The opposite disease phenotypes of the *AtPDF2.2* KD and OE backgrounds support a protective contribution, with the KD phenotype reflecting reduced expression of *AtPDF2.2* and related Class II genes. This protection may involve direct antimicrobial activity, modulation of host responses or both. These findings indicate that the contribution of a defensin to resistance extends beyond its pathogen-induced transcriptional response. The absence of reduced vegetative growth in OE-PDF2.2 under the conditions tested supports further evaluation of *PDF2*-type genes in Brassica.

## Methods

### Plant material and growth conditions

*Arabidopsis thaliana* ecotype Columbia (Col-0) was used as the wild-type background. The hormone-signaling mutants *jar1*, *coi1-16*, *acs1-9*, *ein2-1*, *ein2-1 coi1-16* and *sid2*, and the *AtPDF1.2a* T-DNA insertion line KO-*pdf1.2a*, were obtained from the Nottingham *Arabidopsis* Stock Centre (NASC). Stock accessions, database designations and mutation types are listed in Table S2. The line designated *acs1-9* corresponds to NASC stock N16650 and carries the *acs1-1 acs2-1 acs4-1 acs5-2 acs6-1 acs7-1 acs9-1* allele combination described by Tsuchisaka et al. [41]. KO-*pdf1.2a* corresponds to SALK_063966.49.80.x (NASC N563966), with a T-DNA insertion in the coding region from the SALK collection [42]. Genotyping primers are listed in Table S3.

Seeds were surface-sterilized in 5% (v/v) sodium hypochlorite containing Tween 20, followed by 70% (v/v) ethanol, and washed three times with sterile water. Seeds were plated on half-strength Murashige and Skoog (MS) medium containing 0.8% (w/v) sucrose, 0.1% (w/v) MES and 0.8% (w/v) agar [43]. After stratification at 4 °C for 48 h, seedlings were grown vertically at 22 °C under short-day conditions (8 h light/16 h dark, 100 µmol m⁻² s⁻¹). EMS-derived mutant lines were grown for 30 days at 16 °C under the same photoperiod, together with temperature-matched Col-0 controls, which did not differ from Col-0 grown at 22 °C.

### Sequence retrieval and *in silico* analysis

*AtPDF* family members were retrieved from The *Arabidopsis* Information Resource (TAIR) by sequence similarity search and aligned using ClustalW2. The 50 highest-ranked co-expression partners of each gene were retrieved from ATTED-II [44] using the *Arabidopsis thaliana* ath-u.5 dataset. Gene Ontology and KEGG enrichment analyses were performed with g:Profiler [45] on 13 July 2026 using its multiple-testing correction at a significance threshold of 0.05. Complete co-expression partner lists and enrichment outputs are provided in Tables S4 and S5. Promoter cis-elements within 1.5 kb upstream of the annotated start codon were analyzed using PlantRegMap [46]. Promoter sequences were retrieved from TAIR.

### Pathogen material and inoculation

*Verticillium longisporum* isolate Vl43 was provided by Dr Elke Diederichsen (Freie Universität Berlin, Germany) and maintained on potato dextrose agar. Conidial stocks stored in 22% (v/v) glycerol were centrifuged for 8 min at 8,000 rpm and resuspended in tap water for soil inoculation or sterile Czapek-Dox medium for *in vitro* inoculation to a final concentration of 2 × 10⁶ conidia mL⁻¹. Root inoculation was performed as described by Behrens et al. [13].

The *Sclerotinia sclerotiorum* isolate described by Mei et al. [47] was provided by Professor W. Qian (Southwest University, Chongqing, China). Liquid cultures were prepared as described by Rietz et al. [48] with minor modifications. Five-week-old plants were spray-inoculated with 2 g homogenized mycelium suspended in 50 mL of 10 mM MgSO₄. Mock-treated plants received 10 mM MgSO₄ alone. Plants were maintained under high humidity at 22 °C under short-day conditions.

Disease severity was assessed at 7 dpi using a seven-point scale: 0, no lesion; 1, lesion confined to the inoculation site; 2, leaf lesion; 3, petiole lesion; 4, lesion extending to the stem; 5, stem lesion; and 6, stem death. Disease Severity Index (DSI) was calculated as DSI = Σ(s × nₛ)/(6 × n) × 100, where *nₛ* is the number of plants assigned score *s* and *n* is the total number of plants scored.

For detached-leaf assays, fully expanded leaves from five-week-old plants were placed abaxial side up on moistened filter paper in sealed Petri dishes and inoculated with a mycelial agar plug. Leaves were incubated under the same short-day conditions and photographed at 24 hpi. Lesion diameter was measured in ImageJ from two perpendicular measurements per lesion. Detached-leaf assays were performed with both *AtPDF2.2* and *AtPDF1.2a* transgenic lines.

*Pseudomonas syringae* pv. tomato DC3000 was cultured on NYGA medium containing 50 mg L⁻¹ kanamycin and 100 mg L⁻¹ rifampicin for 2 days at 28 °C. Bacteria were resuspended in 10 mM MgCl₂ and adjusted to OD₆₀₀ = 0.2. Five-week-old plants were spray-inoculated with bacterial suspension or mock solution containing 0.04% (v/v) Silwet Gold. Plants were maintained under high humidity for 2 days and subsequently returned to short-day conditions at 22 °C. Bacterial abundance was quantified by DNA-based qPCR following Ross and Somssich [49].

### RNA isolation and RT-qPCR

Total RNA was extracted from 100 mg plant tissue using TRIzol reagent (Thermo Fisher Scientific). RNA quality and concentration were assessed by agarose gel electrophoresis and NanoVue Plus spectrophotometry. One microgram of total RNA was reverse transcribed using the RevertAid First Strand cDNA Synthesis Kit (Thermo Fisher Scientific) with oligo-dT primers.

RT-qPCR was performed using Maxima SYBR Green/ROX qPCR Master Mix on a CFX96 Touch Real-Time PCR Detection System (Bio-Rad). Cycling conditions were 95 °C for 3 min, followed by 45 cycles of 95 °C for 15 s, 58 °C for 15 s and 72 °C for 20 s, with a final step at 95 °C for 10 s. Relative expression was calculated using the efficiency-corrected method of Pfaffl [50] and normalized to *ACT2* (*AT3G18780*). Biological samples were analyzed with two technical replicates. Primer sequences are listed in Table S1.

Transcript levels were normalized to *ACT2.* Relative expression is shown on a log₂ scale where indicated in the figures. The reference gene used for normalization and the groups used for statistical comparison are distinct: statistical comparisons were made against the relevant wild-type or mock-treated control, or between the groups specified in the figure legend.

### Vector construction and plant transformation

A 1,519-bp genomic fragment immediately upstream of the *AtPDF2.2* start codon was amplified and recombined into pDONR201, followed by transfer into pGWB433 [51] to generate the Prom::*AtPDF2.2*::*GUS* construct.

For *AtPDF2.2* knock-down, a 252-bp fragment spanning the coding region and stop codon was cloned in antisense orientation under the CaMV 35S promoter and transferred into pGWB402 [51]. For overexpression, the full-length coding sequences of *AtPDF2.2* and *AtPDF1.2a* were amplified from Col-0 cDNA and transferred into pGWB402 [51] under the control of the CaMV 35S promoter.

All constructs were verified by sequencing and introduced into *Agrobacterium tumefaciens* GV3101. *Arabidopsis* Col-0 plants were transformed by floral dip [52]. Transformants were selected on half-strength MS medium containing 50 mg L⁻¹ kanamycin. Four independent transformation events were recovered for each construct and screened by RT-qPCR. The three highest-expressing OE events and three lowest-expressing KD events were selected for further analysis. Three homozygous KO-*pdf1.2a* segregant lines were established from the same SALK stock and confirmed by genotyping; these represent the same insertion allele rather than independent insertion events. All lines used in this study are listed in Table S2, and cloning primers are provided in Table S1.

### Genomic DNA isolation and genotyping

Genomic DNA was extracted from frozen plant tissue using CTAB buffer supplemented with β-mercaptoethanol [53]. PCR amplification was performed using 100 ng genomic DNA under the following conditions: 95 °C for 10 min; 30 cycles of 95 °C for 45 s, 60 °C for 30 s and 72 °C for 40 s; followed by 72 °C for 7 min. PCR products were resolved on 1% (w/v) agarose gels.

T-DNA insertions in *jar1*, *ein2-1*, *sid2* and KO-*pdf1.2a* were confirmed using the SALK left-border primer LBb1.3, LP and RP primers. The *COI1* locus in *coi1-16* and *ein2-1 coi1-16* was amplified and sequenced as described by Ellis and Turner [54]. Genotyping primers are listed in Table S3.

### Histochemical GUS analysis

Histochemical detection of β-glucuronidase activity in Prom::*AtPDF2.2*::*GUS* plants was performed according to Jefferson et al. [55]. Seedlings were grown on half-strength MS medium for two weeks before treatment and sampling. Stained tissues were cleared in ethanol and imaged using an AxioVision rel. 4.8 system (Carl Zeiss Microscopy GmbH).

### Quantification of pathogen biomass *in planta*

Genomic DNA was extracted from infected plant material as described above. *V. longisporum* DNA was quantified by real-time PCR using ITS primers OLG70/OLG71 [56] and normalized to an *Arabidopsis* genomic reference amplicon in AT4G26410 [57]. Fungal DNA qPCR cycling conditions were 95 °C for 3 min, followed by 35 cycles of 95 °C for 20 s, 58 °C for 20 s and 72 °C for 30 s. Bacterial abundance was quantified from the Pst DC3000 *oprF* locus following Ross and Somssich [49]. Primer sequences are listed in Table S1.

### Morphological and developmental analysis

For root measurements, approximately 15 seedlings per line were grown vertically on half-strength MS medium containing 0.5% (w/v) sucrose. Plates were photographed at 20 and 30 days after germination (DAG), and primary root length and lateral root number were measured using ImageJ.

For flowering and rosette measurements, plants were grown in soil under short-day conditions. Bolting was recorded daily. Rosette area was measured at approximately 35-40 days after germination using ImageJ to capture developmental differences among the genotypes. Total leaf number was recorded at 35 DAG. Representative rosette photographs were taken at the ages indicated in Fig. S1.

### Statistical analysis

Experiments were performed using a completely randomized design and repeated at least twice with comparable results. Sample sizes are given in the corresponding figure legends. RT-qPCR experiments contained three to four biological replicates per genotype and treatment, each analyzed in two technical replicates. Six infected plants were sampled individually for fungal biomass measurements. For bacterial quantification, each biological replicate consisted of three leaf discs pooled from two plants. Approximately 15 seedlings per line were used for root and rosette measurements.

For experiments combining multiple lines, measurements from individual plants or biological replicates were first averaged within each line. The resulting line means were used as the statistical values (n = 3 lines per genotype), with standard deviation shown as error bars. For OE and KD genotypes, the lines represent independent transformation events. Statistical comparisons were performed using two-tailed Student’s t-tests against the correspondingly treated WT-Col-0 unless otherwise stated in the figure legend.

## Accession numbers

Sequence data for the genes analyzed in this study are available in The *Arabidopsis* Information Resource (TAIR) under the following accession numbers: *AtPDF1.1*, AT1G75830; *AtPDF1.2a*, AT5G44420; *AtPDF1.3*, AT2G26010; *AtPDF1.4*, AT1G19610; *AtPDF1.5*, AT1G55010; *AtPDF2.1*, AT2G02120; *AtPDF2.2*, AT2G02100; *AtPDF2.3*, AT2G02130; *AtPDF2.4*, AT1G61070; *AtPDF2.5*, AT5G63660; *AtPDF2.6*, AT2G02140; Class III (*AtPDF3.1*/LCR80), AT5G38330; Class III (*AtPDF3.2*/LCR59), AT4G30070; *CRT1a*, AT1G56340; *ACT2*, AT3G18780; *LOX3*, AT1G17420; *OPR3*, AT2G06050; *PR1*, AT2G14610; *ETR2*, AT3G23150; *EIN3*, AT3G20770; *AOC3*, AT3G25780; *WRKY70*, AT3G56400; *WRKY53*, AT4G23810; *COI1*, AT2G39940; *JAR1*, AT2G46370; *EIN2*, AT5G03280; *SID2*, AT1G74710.

## Acknowledgements

The author thanks Prof. Daguang Cai (Institute of Phytopathology, Christian-Albrechts-Universität zu Kiel) for supervising this work and Dr Wanzhi Ye for co-supervision.

## Authors’ contributions

Shailja Singh: Investigation, Methodology, Validation, Formal analysis, Writing - original draft, Writing - review & editing.

## Availability of data and materials

All data generated or analyzed during this study are included in this manuscript and its supplementary information files.

## Declarations

### Ethics approval and consent to participate

Not applicable.

### Consent for publication

Not applicable.

## List of abbreviations

AMP: antimicrobial peptide
DAG: days after germination
DSI: Disease Severity Index
dpi: days post-inoculation
ET: ethylene
hpi: hours post-inoculation
JA: jasmonic acid
KD: knock-down
KO: knock-out
OE: overexpression
Pst: Pseudomonas syringae pv. Tomato
SA: salicylic acid
WT: wild type.

## References

1. Parisi K, Shafee TMA, Quimbar P, van der Weerden NL, Bleackley MR, Anderson MA. The evolution, function and mechanisms of action for plant defensins. Semin Cell Dev Biol. 2019;88:107–18.

2. Thomma BPHJ, Cammue BPA, Thevissen K. Plant defensins. Planta. 2002;216:193–202.

3. Liu Y, Hua YP, Chen H, Zhou T, Yue CP, Huang JY. Genome-scale identification of plant defensin (PDF) family genes and molecular characterization of their responses to diverse nutrient stresses in allotetraploid rapeseed. PeerJ. 2021;9:e12007.

4. Penninckx IAMA, Eggermont K, Terras FRG, Thomma BPHJ, De Samblanx GW, Buchala A, et al. Pathogen-induced systemic activation of a plant defensin gene in *Arabidopsis* follows a salicylic acid-independent pathway. Plant Cell. 1996;8:2309–23.

5. Penninckx IAMA, Thomma BPHJ, Buchala A, Métraux JP, Broekaert WF. Concomitant activation of jasmonate and ethylene response pathways is required for induction of a plant defensin gene in *Arabidopsis*. Plant Cell. 1998;10:2103–13.

6. Manners JM, Penninckx IAMA, Vermaere K, Kazan K, Brown RL, Morgan A, et al. The promoter of the plant defensin gene PDF1.2 from *Arabidopsis* is systemically activated by fungal pathogens and responds to methyl jasmonate but not to salicylic acid. Plant Mol Biol. 1998;38:1071–80.

7. Terras FRG, Eggermont K, Kovaleva V, Raikhel NV, Osborn RW, Kester A, et al. Small cysteine-rich antifungal proteins from radish: their role in host defense. Plant Cell. 1995;7:573–88.

8. Vriens K, Peigneur S, De Coninck B, Tytgat J, Cammue BPA, Thevissen K. The antifungal plant defensin *AtPDF2.3* from *Arabidopsis thaliana* blocks potassium channels. Sci Rep. 2016;6:32121.

9. Li R, Wang B, Wu H, Cheng C. Characterization of plant defensin (PDF) genes in banana (*Musa acuminata*) reveals the antifungal ability of *MaPDF2.2* to *Fusarium* wilt pathogens. Horticulturae. 2025;11:513.

10. Inderbitzin P, Davis RM, Bostock RM, Subbarao KV. The ascomycete *Verticillium longisporum* is a hybrid and a plant pathogen with an expanded host range. PLoS One. 2011;6:e18260.

11. Depotter JRL, Deketelaere S, Inderbitzin P, Tiedemann AV, Höfte M, Subbarao KV, et al. *Verticillium longisporum*, the invisible threat to oilseed rape and other brassicaceous plant hosts. Mol Plant Pathol. 2016;17:1004–16.

12. Floerl S, Majcherczyk A, Possienke M, Feussner K, Tappe H, Gatz C, et al. *Verticillium longisporum* infection affects the leaf apoplastic proteome, metabolome, and cell wall properties in *Arabidopsis thaliana*. PLoS One. 2012;7:e31435.

13. Behrens FH, Schenke D, Hossain R, Ye W, Schemmel M, Bergmann T, et al. Suppression of abscisic acid biosynthesis at the early infection stage of *Verticillium longisporum* in oilseed rape (*Brassica napus*). Mol Plant Pathol. 2019;20:1645–61.

14. Hafiz FB, Geistlinger J, Al Mamun A, Schellenberg I, Neumann G, Rozhon W. Tissue-specific hormone signalling and defence gene induction in an *in vitro* assembly of the rapeseed-*Verticillium longisporum* pathosystem. Int J Mol Sci. 2023;24:10489.

15. van Schie CCN, Takken FLW. Susceptibility genes 101: how to be a good host. Annu Rev Phytopathol. 2014;52:551–81.

16. Pröbsting M, Schenke D, Hossain R, Häder C, Thurau T, Wighardt L, et al. Loss of function of CRT1a (calreticulin) reduces plant susceptibility to *Verticillium longisporum* in both *Arabidopsis thaliana* and oilseed rape (*Brassica napus*). Plant Biotechnol J. 2020;18:2328–44.

17. Ye W, Hossain R, Pröbsting M, Ali AAM, Han L, Miao Y, et al. Knock-out of *BnHva22c* reduces the susceptibility of *Brassica napus* to infection with the fungal pathogen *Verticillium longisporum*. Crop J. 2024;12:503-14.

18. Häffner E, Karlovsky P, Diederichsen E. Genetic and environmental control of the *Verticillium* syndrome in *Arabidopsis thaliana*. BMC Plant Biol. 2010;10:235.

19. Harting R, Höfer A, Tran VT, Weinhold LM, Barghahn S, Schlüter R, et al. A 20-kb lineage-specific genomic region tames virulence in pathogenic amphidiploid *Verticillium longisporum*. Mol Plant Pathol. 2021;22:939–53.

20. Ulrich L, Schmitz J, Thurow C, Gatz C. CORONATINE INSENSITIVE 1-mediated repression of immunity-related genes in *Arabidopsis* roots is lifted upon infection with *Verticillium longisporum*. J Exp Bot. 2025;76:2356–72.

21. Hoelscher MP, Forner J, Calderone S, Krämer C, Taylor Z, Loiacono FV, et al. Expression strategies for the efficient synthesis of antimicrobial peptides in plastids. Nat Commun. 2022;13:5856.

22. Yu Z, Xu Y, Zhu L, Zhang L, Liu L, Zhang D, et al. The Brassicaceae-specific secreted peptides, STMPs, function in plant growth and pathogen defense. J Integr Plant Biol. 2020;62:403–20.

23. Tian D, Traw MB, Chen JQ, Kreitman M, Bergelson J. Fitness costs of R-gene-mediated resistance in *Arabidopsis thaliana*. Nature. 2003;423:74–7.

24. Kunkel BN, Brooks DM. Cross talk between signaling pathways in pathogen defense. Curr Opin Plant Biol. 2002;5:325–31.

25. Glazebrook J. Contrasting mechanisms of defense against biotrophic and necrotrophic pathogens. Annu Rev Phytopathol. 2005;43:205–27.

26. Pieterse CMJ, Van der Does D, Zamioudis C, Leon-Reyes A, Van Wees SCM. Hormonal modulation of plant immunity. Annu Rev Cell Dev Biol. 2012;28:489–521.

27. Pré M, Atallah M, Champion A, De Vos M, Pieterse CMJ, Memelink J. The AP2/ERF domain transcription factor ORA59 integrates jasmonic acid and ethylene signals in plant defense. Plant Physiol. 2008;147:1347–57.

28. Zarei A, Körbes AP, Younessi P, Montiel G, Champion A, Memelink J. Two GCC boxes and AP2/ERF-domain transcription factor ORA59 in jasmonate/ethylene-mediated activation of the PDF1.2 promoter in *Arabidopsis*. Plant Mol Biol. 2011;75:321–31.

29. Yao J, Luo JS, Xiao Y, Zhang Z. The plant defensin gene *AtPDF2.1* mediates ammonium metabolism by regulating glutamine synthetase activity in *Arabidopsis thaliana*. BMC Plant Biol. 2019;19:557.

30. Siddique S, Wieczorek K, Szakasits D, Kreil DP, Bohlmann H. The promoter of a plant defensin gene directs specific expression in nematode-induced syncytia in *Arabidopsis* roots. Plant Physiol Biochem. 2011;49:1100–7.

31. Sels J, Mathys J, De Coninck BMA, Cammue BPA, De Bolle MFC. Plant pathogenesis-related (PR) proteins: a focus on PR peptides. Plant Physiol Biochem. 2008;46:941–50.

32. Toruño TY, Stergiopoulos I, Coaker G. Plant-pathogen effectors: cellular probes interfering with plant defenses in spatial and temporal manners. Annu Rev Phytopathol. 2016;54:419–41.

33. Wang M, Weiberg A, Dellota E, Yamane D, Jin H. *Botrytis* small RNA Bc-siR37 suppresses plant defense genes by cross-kingdom RNAi. RNA Biol. 2017;14:421–8.

34. Schenke D, Cai D. Phytohormone crosstalk in the host-Verticillium interaction. Plant Signal Behav. 2020;15:1803567.

35. Häffner E, Karlovsky P, Splivallo R, Traczewska A, Diederichsen E. ERECTA, salicylic acid, abscisic acid, and jasmonic acid modulate quantitative disease resistance of *Arabidopsis thaliana* to *Verticillium longisporum*. BMC Plant Biol. 2014;14:85.

36. Rafiei V, Dixelius C, Tzelepis G. The soilborne fungus *Verticillium longisporum* and its interactions with the brassicaceous hosts. Mol Plant Microbe Interact. 2025;38:497–504.

37. Ke T, Cao H, Huang J, Hu F, Huang J, Dong C, et al. EST-based *in silico* identification and *in vitro* test of antimicrobial peptides in *Brassica napus*. BMC Genomics. 2015;16:653.

38. Dong Y, Wang Y, Tang M, Chen W, Chai Y, Wang W. Bioinformatic analysis of wheat defensin gene family and function verification of candidate genes. Front Plant Sci. 2023;14:1279502.

39. Kaur J, Thokala M, Robert-Seilaniantz A, Zhao P, Peyret H, Berg H, et al. Subcellular targeting of an evolutionarily conserved plant defensin *MtDef4.2* determines the outcome of plant-pathogen interaction in transgenic *Arabidopsis*. Mol Plant Pathol. 2012;13:1032–46.

40. Ding LN, Sun QS, Li JY, Hu YH, Wang J, Tan XL, et al. Molecular switches in plant stress adaptation: the function and engineering of inducible promoters. J Agric Food Chem. 2026;74:13331–48.

41. Tsuchisaka A, Yu G, Jin H, Alonso JM, Ecker JR, Zhang X, et al. A combinatorial interplay among the 1-aminocyclopropane-1-carboxylate isoforms regulates ethylene biosynthesis in *Arabidopsis thaliana*. Genetics. 2009;183:979–1003.

42. Alonso JM, Stepanova AN, Leisse TJ, Kim CJ, Chen H, Shinn P, et al. Genome-wide insertional mutagenesis of *Arabidopsis thaliana*. Science. 2003;301:653–7.

43. Murashige T, Skoog F. A revised medium for rapid growth and bio assays with tobacco tissue cultures. Physiol Plant. 1962;15:473–97.

44. Obayashi T, Hibara H, Kagaya Y, Aoki Y, Kinoshita K. ATTED-II v11: a plant gene coexpression database using a sample balancing technique. Plant Cell Physiol. 2022;63:869–81.

45. Kolberg L, Raudvere U, Kuzmin I, Adler P, Vilo J, Peterson H. g:Profiler-interoperable web service for functional enrichment analysis and gene identifier mapping (2023 update). Nucleic Acids Res. 2023;51:W207–12.

46. Tian F, Yang DC, Meng YQ, Jin J, Gao G. PlantRegMap: charting functional regulatory maps in plants. Nucleic Acids Res. 2020;48:D1104–13.

47. Mei J, Wei D, Disi JO, Ding Y, Liu Y, Qian W. Screening resistance against *Sclerotinia sclerotiorum* in Brassica crops with use of detached stem assay under controlled environment. Eur J Plant Pathol. 2012;134:599–604.

48. Rietz S, Bernsdorff FEM, Cai D. Members of the germin-like protein family in *Brassica napus* are candidates for the initiation of an oxidative burst that impedes pathogenesis of *Sclerotinia sclerotiorum*. J Exp Bot. 2012;63:5507–19.

49. Ross A, Somssich IE. A DNA-based real-time PCR assay for robust growth quantification of the bacterial pathogen *Pseudomonas syringae* pv. tomato in *Arabidopsis thaliana*. Plant Methods. 2016;12:48.

50. Pfaffl MW. A new mathematical model for relative quantification in real-time RT-PCR. Nucleic Acids Res. 2001;29:e45.

51. Nakagawa T, Suzuki T, Murata S, Nakamura S, Hino T, Maeo K, et al. Improved Gateway binary vectors: high-performance vectors for creation of fusion constructs in transgenic analysis of plants. Biosci Biotechnol Biochem. 2007;71:2095–100.

52. Clough SJ, Bent AF. Floral dip: a simplified method for Agrobacterium-mediated transformation of *Arabidopsis thaliana*. Plant J. 1998;16:735–43.

53. Rogers SO, Bendich AJ. Extraction of DNA from milligram amounts of fresh, herbarium and mummified plant tissues. Plant Mol Biol. 1985;5:69–76.

54. Ellis C, Turner JG. A conditionally fertile coi1 allele indicates cross-talk between plant hormone signalling pathways in *Arabidopsis thaliana* seeds and young seedlings. Planta. 2002;215:549–56.

55. Jefferson RA, Kavanagh TA, Bevan MW. GUS fusions: beta-glucuronidase as a sensitive and versatile gene fusion marker in higher plants. EMBO J. 1987;6:3901–7.

56. Eynck C, Koopmann B, Grunewaldt-Stoecker G, Karlovsky P, von Tiedemann A. Differential interactions of *Verticillium longisporum* and *V. dahliae* with *Brassica napus* detected with molecular and histological techniques. Eur J Plant Pathol. 2007;118:259–74.

57. Czechowski T, Stitt M, Altmann T, Udvardi MK, Scheible WR. Genome-wide identification and testing of superior reference genes for transcript normalization in *Arabidopsis*. Plant Physiol. 2005;139:5–17.

