## SUPPORTING INFORMATION for "*PDF2.2* links constitutive defensin expression to disease resistance in *Arabidopsis*"

### Supplementary figures

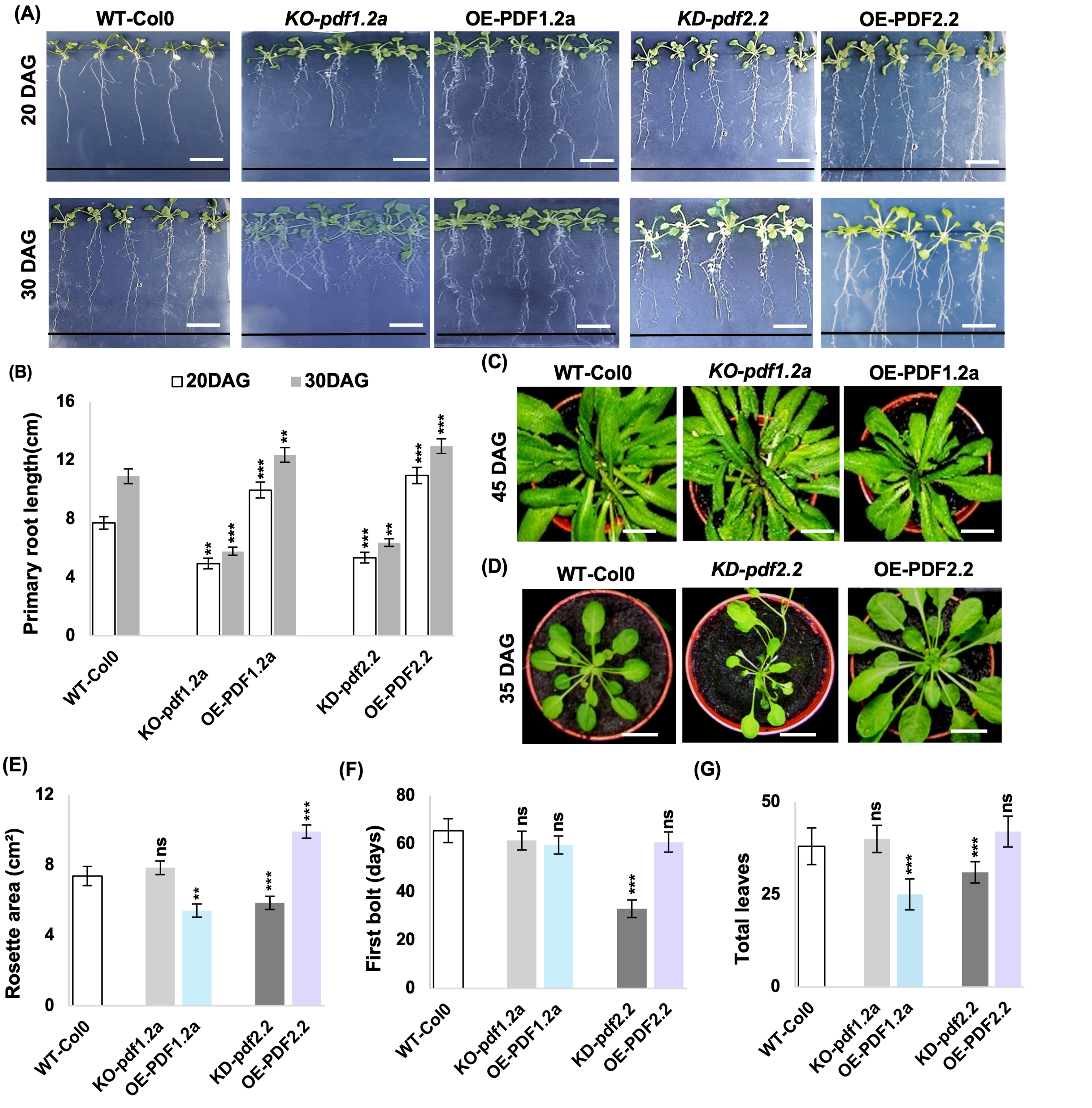


**Figure S1.** Developmental phenotypes under non-infected conditions. (A) Root systems at 20 and 30 days after germination (DAG). Bars = 1 cm. (B) Primary root length (cm) at both time points. (C) Rosettes of WT-Col-0, KO-*pdf1.2a* and OE-PDF1.2a at 45 DAG. Bars = 1 cm. (D) Rosettes of WT-Col-0, KD-*pdf2.2* and OE-PDF2.2 at 35 DAG. Bars = 1 cm. (E) Rosette area ($cm^2$) of non-infected plants at approximately 35–40 DAG. (F) Days to first bolt. (G) Total leaf number, counted at 35 DAG. (B, E, F, G) Data are mean ± SD of three lines (n = 3); each line value represents the mean of the biological replicates within that line. Asterisks indicate significant differences from WT-Col-0 (Student's t-test): **P < 0.05, **P < 0.01, ***P < 0.005; ns, not significant.*

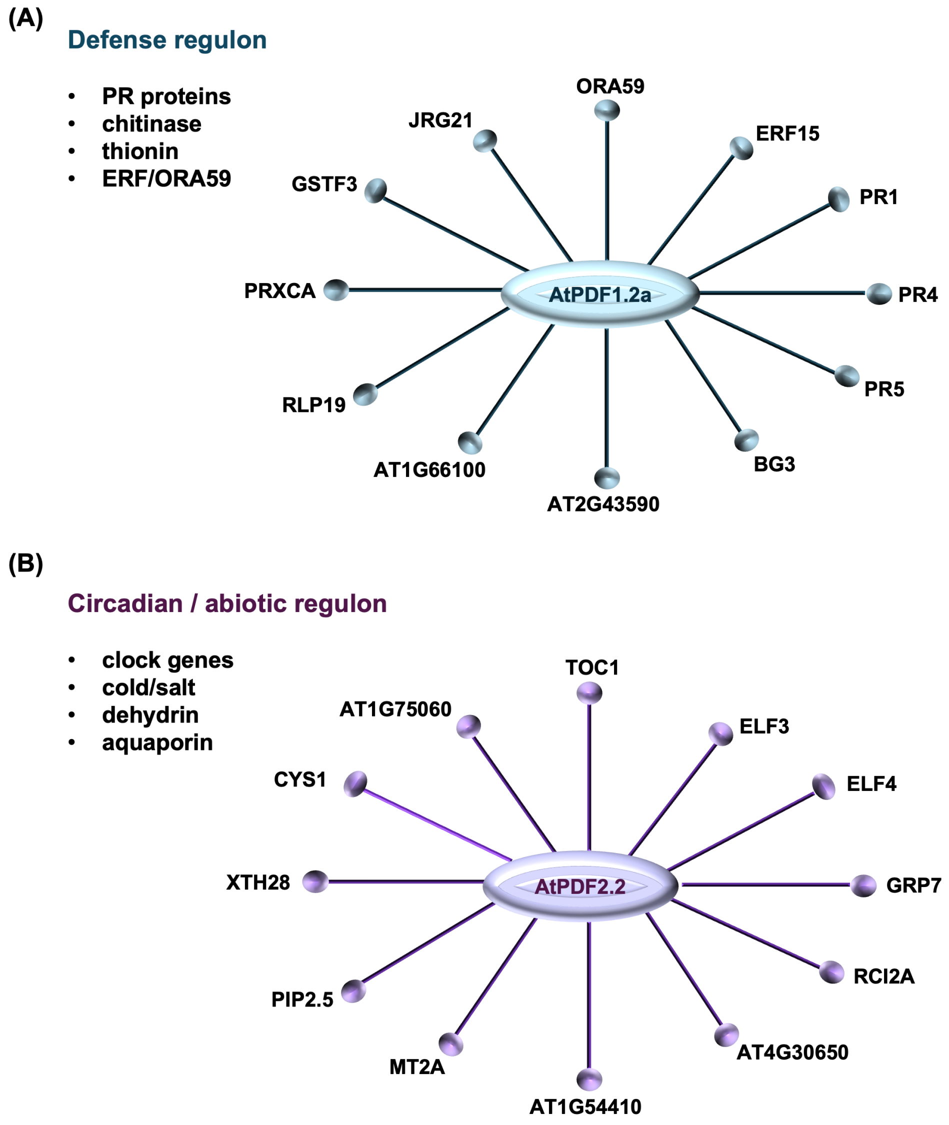


**Figure S2.** *AtPDF1.2a* and *AtPDF2.2* occupy distinct co-expression neighborhoods. ATTED-II co-expression networks. (A) *AtPDF1.2a* is associated with defense-related genes, including PR proteins, chitinase, β-1,3-glucanase and components of the ERF/ORA59 network. (B) *AtPDF2.2* is preferentially associated with circadian- and abiotic-stress-related genes, including TOC1, ELF3, ELF4, GRP7, RCI2A and PIP2;5. These networks show co-expression associations.

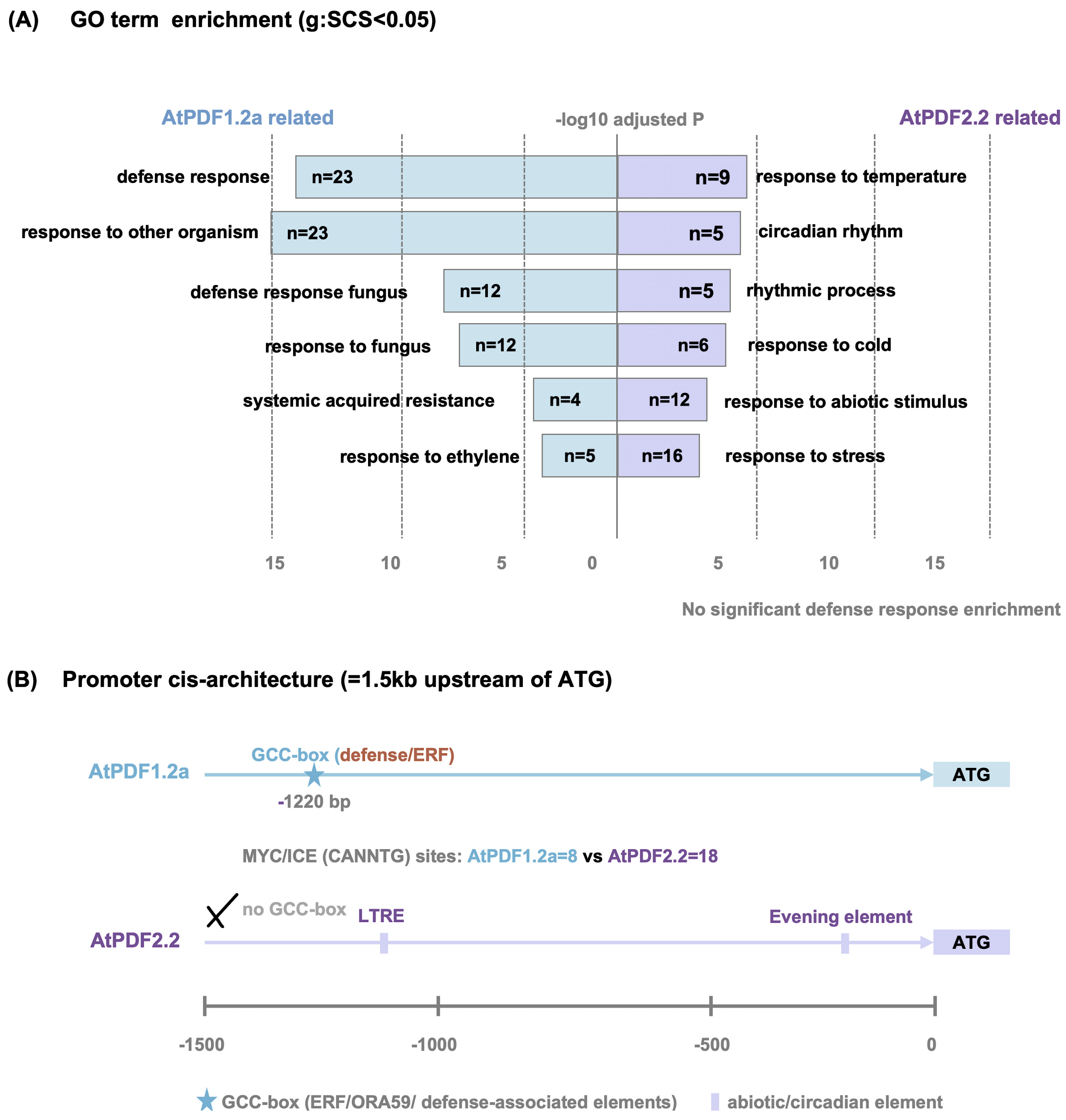


**Figure S3.** GO-term enrichment and promoter architecture of *AtPDF1.2a* and *AtPDF2.2*. (A) GO-term enrichment of the co-expression partners (g:Profiler, g:SCS < 0.05). *AtPDF1.2a* partners are enriched for defense- and fungus-associated processes, whereas *AtPDF2.2* partners are enriched for temperature, circadian and abiotic-stress responses. (B) Promoter cis-architecture approximately 1.5 kb upstream of the ATG. The *AtPDF1.2a* promoter contains a GCC-box (ERF/ORA59 element) at approximately -1,220 bp. The analyzed *AtPDF2.2* promoter lacks a GCC-box and contains an LTRE, an Evening Element and 18 MYC/ICE (CANNTG) sites compared with eight in *AtPDF1.2a*. These *in silico* features indicate distinct promoter architectures; their functional contribution was not investigated.

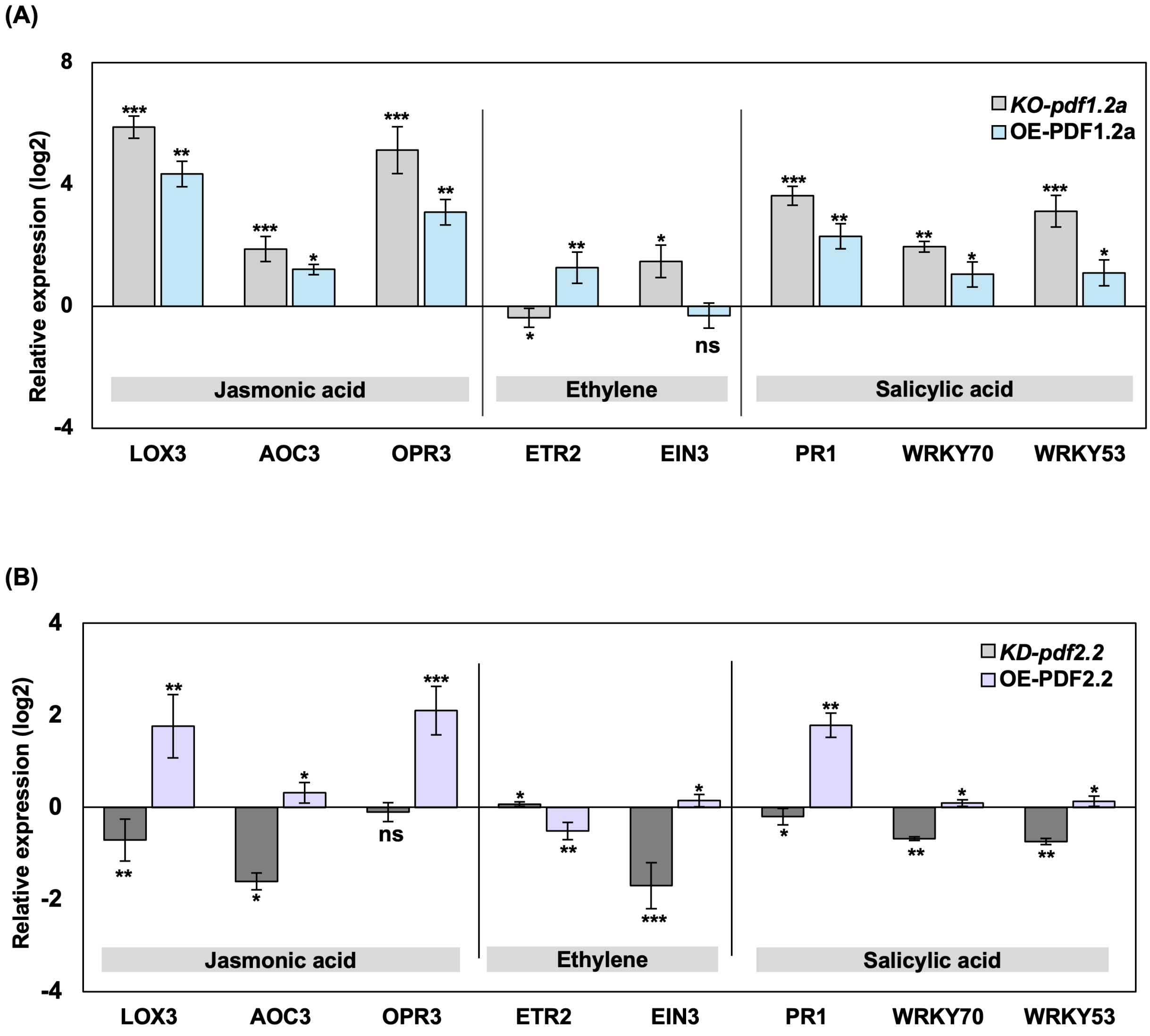


**Figure S4.** Defense-hormone marker gene expression in the transgenic lines. Transcript levels of the jasmonic acid biosynthesis genes LOX3, AOC3 and OPR3, the ethylene-signaling regulators ETR2 and EIN3, and the salicylic acid-associated genes PR1, WRKY70 and WRKY53 in Vl43-infected transgenic lines, determined by RT-qPCR, normalized to *ACT2* and expressed as $log_2$ relative expression. (A) KO-*pdf1.2a* and OE-PDF1.2a. (B) KD-*pdf2.2* and OE-PDF2.2. Data are mean ± SD of three biological replicates. Asterisks indicate significant differences from infected WT-Col-0 (Student's t-test): **P < 0.05, **P < 0.01, ***P < 0.005; ns, not significant.*

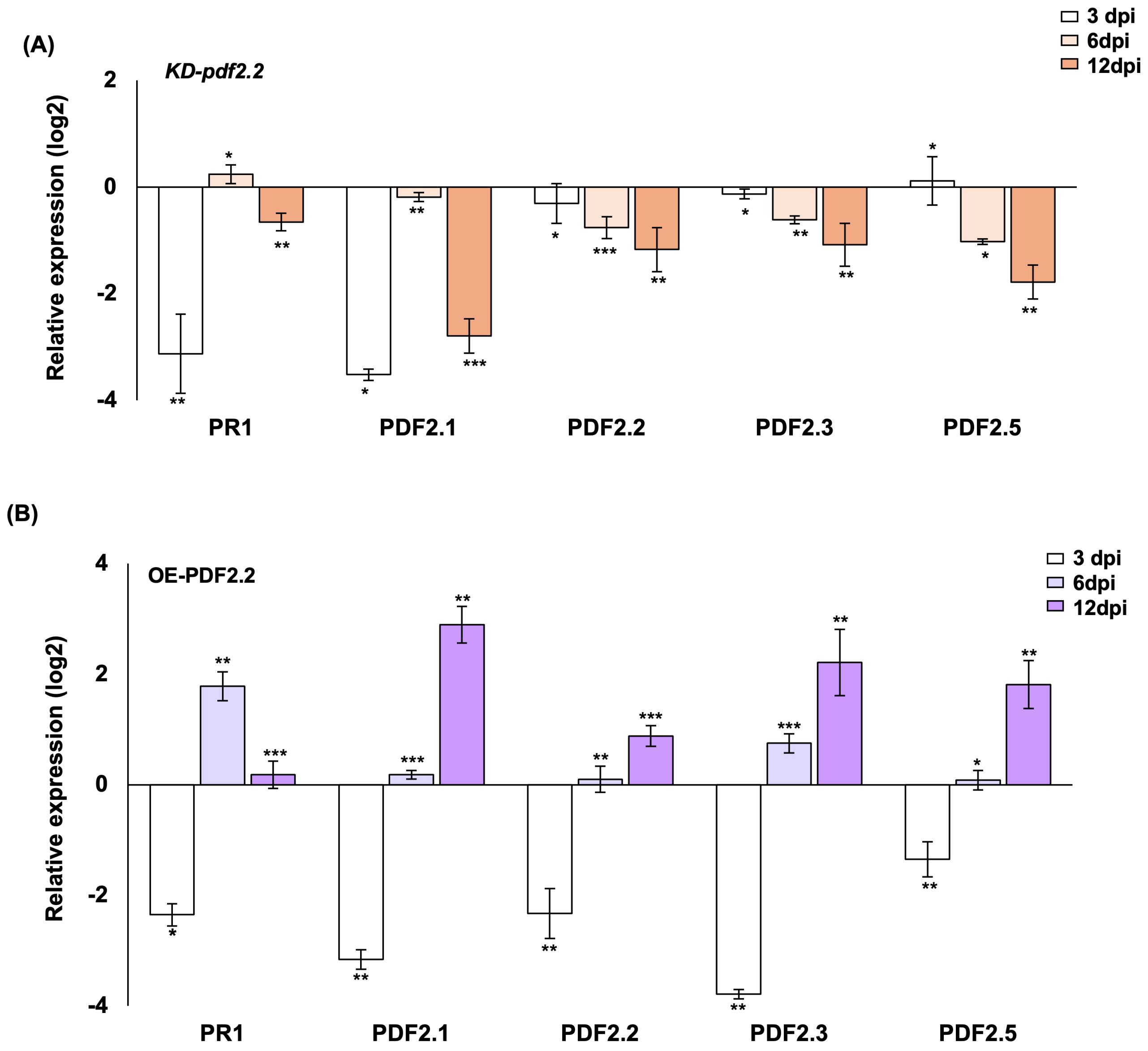


**Figure S5.** Class II defensin expression in *AtPDF2.2* transgenic lines during *Verticillium longisporum* infection. Transcript levels of PR1 and the Class II defensins *AtPDF2.1*, *AtPDF2.2*, *AtPDF2.3* and *AtPDF2.5* in roots of (A) KD-*pdf2.2* and (B) OE-PDF2.2 lines at 3, 6 and 12 dpi with Vl43, determined by RT-qPCR, normalized to *ACT2* and expressed as $log_2$ relative expression. Data are mean ± SD of three to four biological replicates. Asterisks indicate significant differences from infected WT-Col-0 (Student's t-test*): *P < 0.05, **P < 0.01, ***P < 0.005; ns, not significant.*

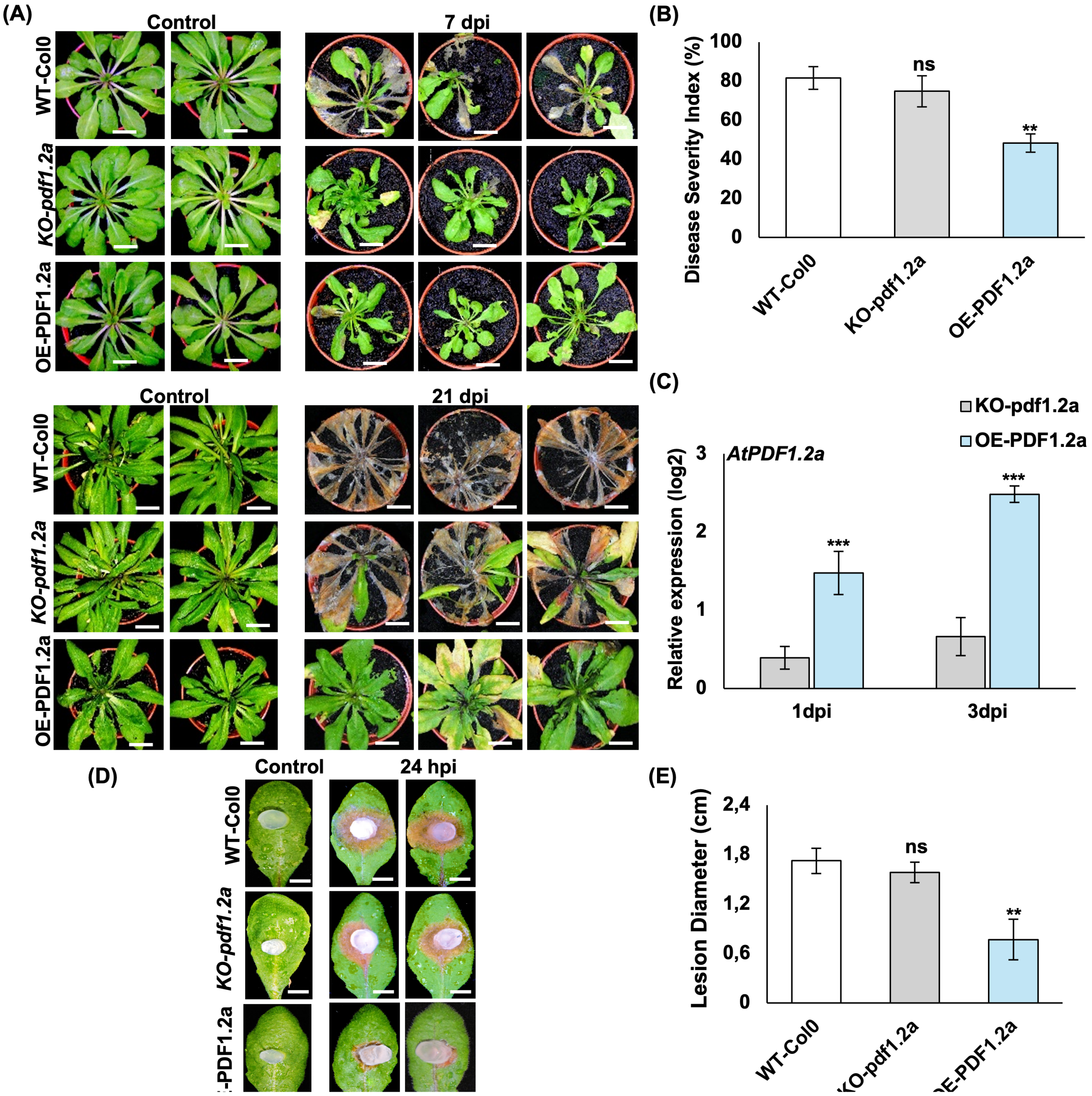


**Figure S6.** *AtPDF1.2a* transgenic lines under *Sclerotinia sclerotiorum* infection. (A) Whole-plant phenotypes at 7 and 21 dpi and in the control. Bars = 1 cm. (B) Disease Severity Index (%) scored at 7 dpi on the 0-6 scale defined in Methods. (C) Transcript levels of *AtPDF1.2a* at 1 and 3 dpi, determined by RT-qPCR, normalized to *ACT2* and expressed as $log_2$ relative expression. (D) Detached-leaf assay at 24 hpi. Bars = 1 cm. (E) Lesion diameter (cm). (B, E) Data are mean ± SD across three lines (n = 3). (C) Data are mean ± SD of three biological replicates. Asterisks indicate significant differences from the correspondingly treated WT-Col-0 (Student's t-test): **P < 0.05, **P < 0.01, ***P < 0.005; ns, not significant.*

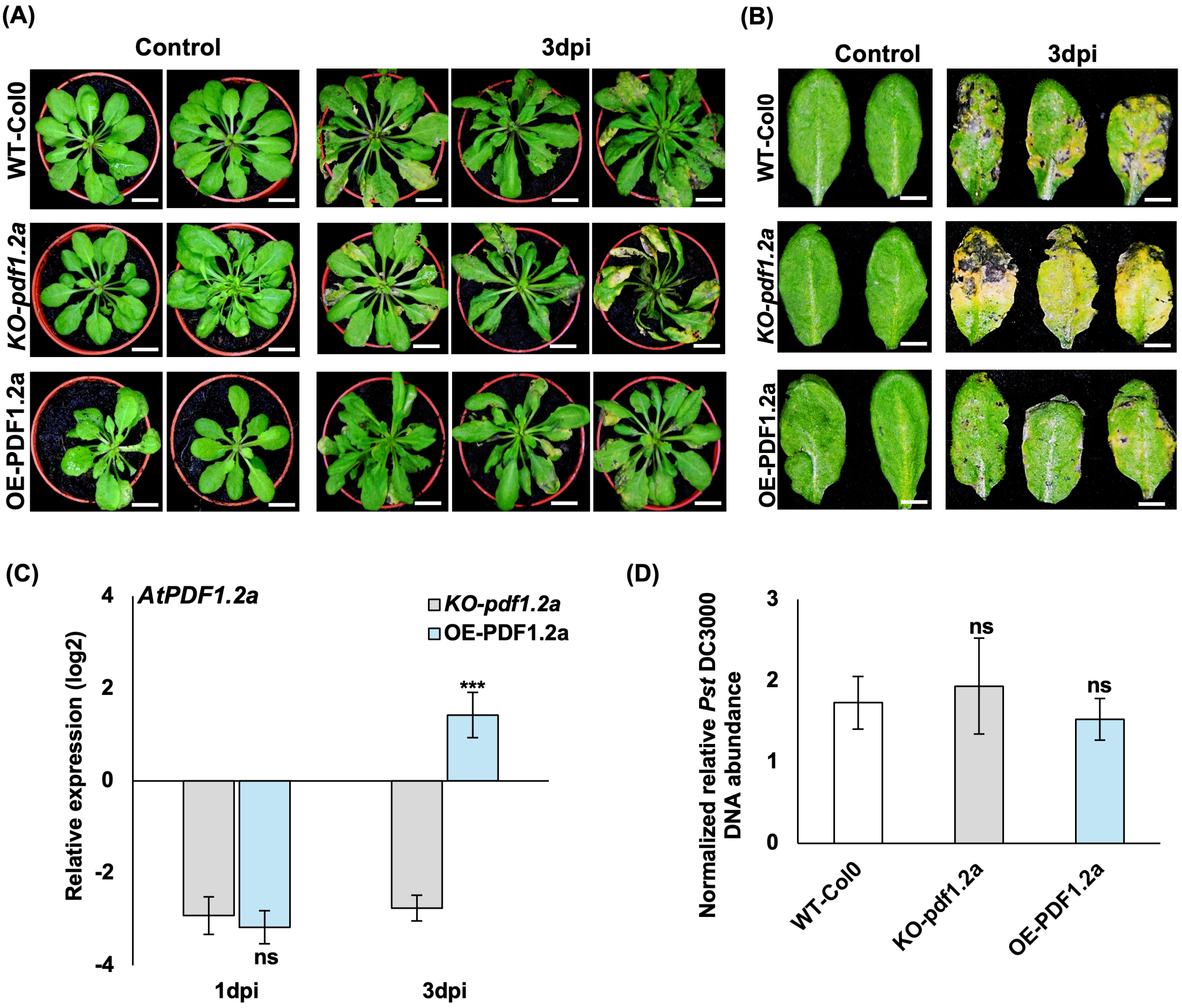


**Figure S7.** *AtPDF1.2a* transgenic lines under Pst DC3000 infection. (A) Rosette symptoms under control conditions and at 3 dpi. Bars = 1 cm. (B) Detached-leaf symptoms of the same genotypes. Bars = 1 cm. (C) Transcript levels of *AtPDF1.2a* at 1 and 3 dpi, determined by RT-qPCR, normalized to *ACT2* and expressed as $log_2$ relative expression. Data are mean ± SD of three to four biological replicates. Asterisks in C indicate significant differences between OE-PDF1.2a and KO-*pdf1.2a* rather than from wild type (Student's t-test): *P < 0.05, **P < 0.01, ***P < 0.005; ns, not significant. (D) Normalized relative Pst DC3000 DNA abundance. Data are mean ± SD of three lines (n = 3); each line value represents the mean of the biological replicates within that line. Asterisks indicate significant differences from the correspondingly treated WT-Col-0 (Student's t-test): **P < 0.05, **P < 0.01, ***P < 0.005; ns, not significant.*

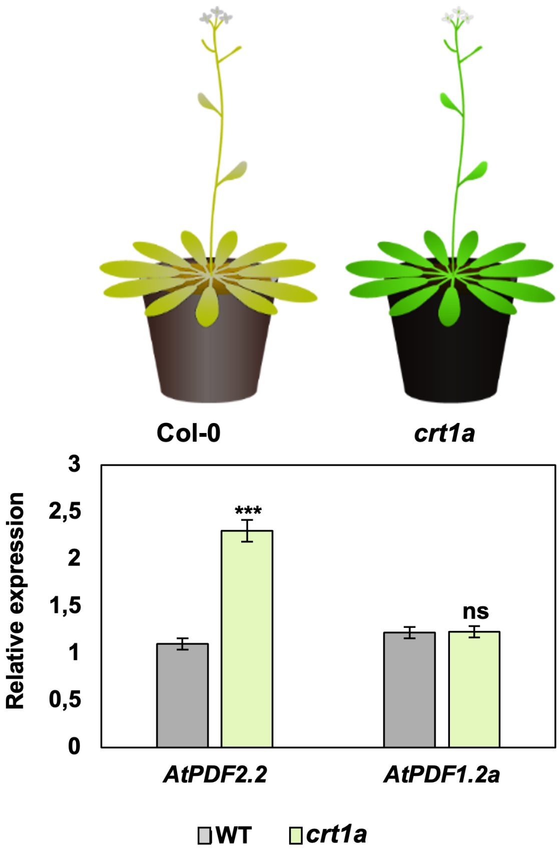


**Figure S8.** *AtPDF2.2* and *AtPDF1.2a* expression in the *crt1a* mutant. Schematic rosette phenotypes of Col-0 and *crt1a*, and transcript levels of *AtPDF2.2* and *AtPDF1.2a* in Vl43-infected WT-Col-0 and *crt1a*, determined by RT-qPCR relative to *ACT2*. *crt1a* is a T-DNA knock-out of the susceptibility factor CRT1a [16]. Data are mean ± SD of three to four biological replicates. Asterisks indicate significant differences from infected WT-Col-0 (Student's t-test): **P < 0.05, **P < 0.01, ***P < 0.005; ns, not significant.*

### Supplementary tables

**Table S1 |** Primers used for gene expression, construct/transgene analysis and pathogen/plant biomass quantification.

| Gene/target | Locus/accession | Forward primer (5′→3′) | Reverse primer (5′→3′) |
|---|---|---|---|
| AtPDF1.1 | AT1G75830 | CACAGAAGTTGTGCGAGAGG | GCAAGATCCATGTCGTGCTT |
| AtPDF1.2a | AT5G44420 | CGCTGCTCTTGTTCTCTTTGC | TCCATGTTTGGCTCCTTCAA |
| OE-PDF1.2a (transgene) | AT5G44420 | ACATGGTCAGGGGTTTGCGGAAAC | ATGGAAGTAGTCTTACGTCGTACA |
| AtPDF1.3 | AT2G26010 | CCTCTTCGCTGCTCTTGTTC | ACTTGTGAGCTGGGAAGACA |
| AtPDF1.4 | AT1G19610 | AGTCATGGCTTCTTCTTACACACT | TGAAGTAGCAGAAACATGCGAA |
| AtPDF1.5 | AT1G55010 | TGAAGCACCGACAATTGTGAA | CCACCAGCGCAATATCCATC |
| AtPDF2.1 | AT2G02120 | GATCGACGATTGAGTCATGC | CTCAGCCGTTCTTTTCTTGG |
| AtPDF2.2 | AT2G02100 | GCACCAGACATTGCTGATCT | AACCGGAAACACACAAATCC |
| OE-PDF2.2 (transgene) | AT2G02100 | TCTCTTGCAATGAAGCTCTCTATG | ATGGGTAGATCAGCAATGTCTGGT |
| KD-pdf2.2 (transgene) | AT2G02100 | TCTCTTGCAATGAAGCTCTCTATG | ATGGGTAGATCAGCAATGTCTGGT |
| Prom::AtPDF2.2 | AT2G02100 | CTTCTACGCATGTAAATTATAACC | CAAGTGTGTTGCTGCTAGTGA |
| AtPDF2.3 | AT2G02130 | CACACACAACTGTGCAAACG | CGGAAACACACAAACCAATG |
| AtPDF2.4 | AT1G61070 | ACAGTGATGGGTCTGGTCAC | AAGGTCTGGTGCACAAACAG |
| AtPDF2.5 | AT5G63660 | TGGAGAACAAGTTTTTCGCTGC | AAACTAGTTAGCAAAGGCGGG |
| AtPDF2.6 | AT2G02140 | ATGTCGGTCATGCTCCTCTT | TTTGGCACAGCTCTGTGAGT |
| AtPDF3.1 (LCR80) | AT5G38330 | CCGGTTGGAATGTTGCTGTG | AATTGCCAAGCCCACCATTG |
| AtPDF3.2 (LCR59) | AT4G30070 | TGATCGTAGTGTTGGGGTGC | CATATACCAGCCCCACCGTC |
| ACT2 | AT3G18780 | ACCTTGCTGGACGTGACCTTACTGAT | GTTGTCTCGTGGATTCCAGCAGCTT |
| PR1 | AT2G14610 | TCAGGTTGTTTGGAGAAAGTC | CATTAGTAAGGCTTCTCGTTCACA |
| ETR2 | AT3G23150 | CGTTGGAATTTCACAGGTCGATGAG | CGTCTTCGCAGTTACATCGTGGA |
| EIN3 | AT3G20770 | TGGTGGCCTCAACTTGGTTT | CGAGTTTCCTGCTGGGACTT |
| AOC3 | AT3G25780 | GAATTGGGGCAAGAAGACCGAAA | TTGCCGAGTTTAAGAACTGCTGGA |
| WRKY70 | AT3G56400 | CCGCCGTTGAGGGATCT | GGACGAACCATGATGACGAT |
| WRKY53 | AT4G23810 | GCACGACTTAGAGAAGCTC | ACTCCTTGGGAATTTGGCG |
| LOX3 | AT1G17420 | GCCGATCTAATTCGCAGAGGA | AAAAGACCGTCGTTGGCGTA |
| OPR3 | AT2G06050 | TGGTTGGCATGCTCAATAAG | GCCTTCCAGACTCTGTTTGC |
| Plant genomic reference | AT4G26410 | GAGCTGAAGTGGCTTCCATGAC | GGTCCGACATACCCATGATCC |
| GUS | EG11055 | GTTGACTGGCAGGTGGTGG | GGTAGATATCACACTCTGTCTGG |
| V. longisporum ITS | OLG70/71 | CAGCGAAACGCGATATGTAG | GGCTTGTAGGGGGTTTAGA |
| Pst DC3000 biomass (oprF) | oprF | AACTGAAAAACACCTTGGGC | CCTGGGTTGTTGAAGTGGTA |

*Accession numbers beginning with AT are Arabidopsis loci. AT4G26410 is the Arabidopsis genomic reference used for pathogen DNA normalization [57].

**Table S2 |** Arabidopsis mutant and transgenic lines used in this study.

| Line | Database designation / construct | Origin | Stock or event ID | Feature |
|---|---|---|---|---|
| jar1 | SALK_059774C | NASC | N658259 | T-DNA insertion in exon |
| coi1-16 | coi1-16 | NASC | N67817 | EMS-derived point mutation |
| ein2-1 | SALK_086500C | NASC | N654757 | T-DNA insertion in exon |
| ein2-1 coi1-16 | ein2-1; coi1-16 | NASC | N67818 | EMS-derived double mutant |
| acs1-9 | acs1-1 acs2-1 acs4-1 acs5-2 acs6-1 acs7-1 acs9-1 | NASC | N16650 | Higher-order T-DNA insertion mutant [41] |
| sid2 | SALK_088254 | NASC | N65675 | T-DNA insertion in exon |
| KO-pdf1.2a | SALK_063966.49.80.x | NASC | N563966 | T-DNA insertion in exon |
| OE-PDF2.2 | 35S::AtPDF2.2 (pGWB402) | This study | Events #1, #7, #8, #12 | Sense-orientation overexpression |
| KD-pdf2.2 | 35S::AtPDF2.2 antisense (pGWB402) | This study | Events #1, #4, #7, #8 | Antisense-mediated knockdown |
| OE-PDF1.2a | 35S::AtPDF1.2a (pGWB402) | This study | Events #1, #2, #3, #7 | Sense-orientation overexpression |
| Prom::AtPDF2.2::GUS | 1,519 bp promoter in pGWB433 | This study | Events #13, #21, #22 | Transcriptional GUS reporter |

*All lines are in the Columbia-0 (Col-0) background. Mutant lines were obtained from the Nottingham Arabidopsis Stock Centre (NASC).
*The line designated acs1-9 throughout the figures and text is NASC stock N16650, which carries the seven-member allele combination listed above. The single-allele designation is retained for consistency with the figure panels.

**Table S3 |** Primers used for genotyping of mutant lines.

| Primer name | Sequence (5′→3′) | Description |
|---|---|---|
| LBb1.3 | ATTTTGCCGATTTCGGAAC | Standard SALK T-DNA left-border primer |
| AtPDF1.2a_LP | ACGTTGATCTTTGGCTTATTTCTTC | SALK_063966.49.80.x LP primer for AtPDF1.2a |
| AtPDF1.2a_RP | CATTGCCGGTGCGTCTGCATAT | SALK_063966.49.80.x RP primer for AtPDF1.2a |
| JAR1_LP | ACGCTCATCAAGTCCAGAAACA | SALK_059774C LP primer for JAR1 |
| JAR1_RP | GGAACAGGGTGACCAGTGAG | SALK_059774C RP primer for JAR1 |
| EIN2_LP | ACAAAGAGGTTCGTCACCGT | SALK_086500C LP primer for EIN2 |
| EIN2_RP | TGACGGATCCCAGTATCCACT | SALK_086500C RP primer for EIN2 |
| SID2_LP | TCTCCCGCAAGGTCTTTTT | SALK_088254 LP primer for SID2 |
| SID2_RP | TCAGAGACGGCGGAGATTAG | SALK_088254 RP primer for SID2 |
| ACS1_sense | TCGACTAAGTTTCCGACGAC | Sense primer for ACS1 (AT3G61510) in the acs1-9 line [41] |
| ACS1_antisense | TGTCGAAATTGAATTATTCCA | Antisense primer for ACS1 (AT3G61510) in the acs1-9 line [41] |
| ACS2_LP | GCAGTTTTGTCGAAGATAGCG | T-DNA LP primer for acs2-1 [41] |
| ACS2_RP | ATCTGTACCTGATTCTCTGCAA | T-DNA RP primer for acs2-1 [41] |
| COI1_F | ACCACAAATAAAGTTTCAGATGCCA | Flanking primer for COI1 sequencing [54] |
| COI1_R | CCCCGAAAACTATGTCGGCT | Flanking primer for COI1 sequencing [54] |

*T-DNA insertion lines were genotyped with locus-specific left (LP) and right (RP) primers together with the SALK T-DNA left-border primer LBb1.3. EMS-derived lines were verified by amplification and sequencing of the target locus.

**Table S4 |** ATTED-II co-expression partners of *AtPDF1.2a* and *AtPDF2.2*.

### *AtPDF1.2a (AT5G44420)*

| Rank | Gene | AGI locus | Annotation |
|---|---|---|---|
| 1 | PDF1.2b | AT2G26020 | plant defensin 1.2b |
| 2 | PDF1.2c | AT5G44430 | plant defensin 1.2C |
| 3 | PDF1.3 | AT2G26010 | plant defensin 1.3 |
| 4 | AACT1 | AT5G61160 | anthocyanin 5-aromatic acyltransferase 1 |
| 5 | PR4 | AT3G04720 | pathogenesis-related 4 |
| 6 | AT3G23550 | AT3G23550 | MATE efflux family protein |
| 7 | ORA59 | AT1G06160 | octadecanoid-responsive AP2/ERF 59 |

| Rank | Gene | AGI locus | Annotation |
|---|---|---|---|
| 8 | DIN11 | AT3G49620 | 2-oxoglutarate (2OG) and Fe(II)-dependent oxygenase superfamily protein |
| 9 | NATA1 | AT2G39030 | Acyl-CoA N-acyltransferases (NAT) superfamily protein |
| 10 | AT4G16260 | AT4G16260 | Glycosyl hydrolase superfamily protein |
| 11 | AT1G15125 | AT1G15125 | S-adenosyl-L-methionine-dependent methyltransferases superfamily protein |
| 12 | CYP71B22 | AT3G26200 | cytochrome P450, family 71, subfamily B, polypeptide 22 |
| 13 | AT4G24350 | AT4G24350 | Phosphorylase superfamily protein |
| 14 | AT4G27300 | AT4G27300 | S-locus lectin protein kinase family protein |
| 15 | AT3G16530 | AT3G16530 | Legume lectin family protein |
| 16 | AT2G43590 | AT2G43590 | Chitinase family protein |
| 17 | AT3G15356 | AT3G15356 | Legume lectin family protein |
| 18 | UGT71C3 | AT1G07260 | UDP-glucosyl transferase 71C3 |
| 19 | AIG2 | AT3G28930 | AIG2-like (avirulence induced gene) family protein |
| 20 | AT1G78460 | AT1G78460 | SOUL heme-binding family protein |
| 21 | AT5G58390 | AT5G58390 | Peroxidase superfamily protein |
| 22 | AT5G42530 | AT5G42530 | uncharacterized protein |
| 23 | GLR5 | AT2G32400 | glutamate receptor 5 |
| 24 | AT5G11610 | AT5G11610 | Exostosin family protein |
| 25 | ERF15 | AT2G31230 | ethylene-responsive element binding factor 15 |
| 26 | CLH1 | AT1G19670 | chlorophyllase 1 |
| 27 | CYP71B36 | AT3G26320 | cytochrome P450, family 71, subfamily B, polypeptide 36 |
| 28 | RLP19 | AT2G15080 | receptor like protein 19 |
| 29 | BG3 | AT3G57240 | beta-1,3-glucanase 3 |
| 30 | AT2G15042 | AT2G15042 | Leucine-rich repeat (LRR) family protein |
| 31 | SOS4 | AT5G37850 | pfkB-like carbohydrate kinase family protein |
| 32 | AT3G16670 | AT3G16670 | Pollen Ole e 1 allergen and 12xtension family protein |
| 33 | AT4G11290 | AT4G11290 | Peroxidase superfamily protein |
| 34 | LSH10 | AT2G42610 | LIGHT-DEPENDENT SHORT HYPOCOTYLS-like protein (DUF640) |
| 35 | GSTF3 | AT2G02930 | glutathione S-transferase F3 |
| 36 | PR1 | AT2G14610 | pathogenesis-related protein 1 |
| 37 | JAL23 | AT2G39330 | jacalin-related lectin 23 |
| 38 | PRXCA | AT3G49110 | peroxidase CA |

| Rank | Gene | AGI locus | Annotation |
|---|---|---|---|
| 39 | ATMDAR2 | AT5G03630 | Pyridine nucleotide-disulfide oxidoreductase family protein |
| 40 | PR5 | AT1G75040 | pathogenesis-related protein 5 |
| 41 | CYP81D1 | AT5G36220 | cytochrome p450 81d1 |
| 42 | JRG21 | AT3G55970 | jasmonate-regulated gene 21 |
| 43 | AT1G66100 | AT1G66100 | Plant thionin |
| 44 | RAP2.9 | AT4G06746 | uncharacterized protein |
| 45 | BGLU18 | AT1G52400 | beta glucosidase 18 |
| 46 | PME3 | AT3G14310 | pectin methylesterase 3 |
| 47 | AT5G41761 | AT5G41761 | uncharacterized protein |
| 48 | PLA2A | AT2G26560 | phospholipase A 2A |
| 49 | AT4G12490 | AT4G12490 | Bifunctional inhibitor/lipid-transfer protein/seed storage 2S albumin superfamily protein |
| 50 | SD1-13 | AT1G11350 | S-domain-1 13 |

***AtPDF2.2 (AT2G02100)***

| Rank | Gene | AGI locus | Annotation |
|---|---|---|---|
| 1 | PDF2.1 | AT2G02120 | Scorpion toxin-like knottin superfamily protein |
| 2 | LCR68 | AT2G02130 | low-molecular-weight cysteine-rich 68 |
| 3 | FP6 | AT4G38580 | farnesylated protein 6 |
| 4 | AT4G30650 | AT4G30650 | Low temperature and salt responsive protein family |
| 5 | ACA8 | AT5G57110 | autoinhibited Ca2+ -ATPase |
| 6 | AT1G68220 | AT1G68220 | aerobic coproporphyrinogen-III oxidase (DUF1218) |
| 7 | AT5G39410 | AT5G39410 | Saccharopine dehydrogenase |
| 8 | AMI1 | AT1G08980 | amidase 1 |
| 9 | E13L3 | AT5G08000 | glucan endo-1,3-beta-glucosidase-like protein 3 |
| 10 | AT2G32280 | AT2G32280 | GPI inositol-deacylase C, putative (DUF1218) |
| 11 | RCI2A | AT3G05880 | Low temperature and salt responsive protein family |
| 12 | GRP7 | AT2G21660 | cold, circadian rhythm, and rna binding 2 |
| 13 | TOC1 | AT5G61380 | CCT motif -containing response regulator protein |
| 14 | AT1G13930 | AT1G13930 | oleosin-B3-like protein |
| 15 | PMR5 | AT5G58600 | Pmr5/Cas1p GDSL/SGNH-like acyl-esterase family protein (DUF828) |

| Rank | Gene | AGI locus | Annotation |
|---|---|---|---|
| 16 | AT1G25320 | AT1G25320 | Leucine-rich repeat protein kinase family protein |
| 17 | AT1G53035 | AT1G53035 | uncharacterized protein |
| 18 | PIP2;5 | AT3G54820 | plasma membrane intrinsic protein 2;5 |
| 19 | AT4G26670 | AT4G26670 | Mitochondrial import inner membrane translocase subunit Tim17/Tim22/Tim23 family protein |
| 20 | ELF4 | AT2G40080 | EARLY FLOWERING-like protein (DUF1313) |
| 21 | SCPL49 | AT3G10410 | SERINE CARBOXYPEPTIDASE-LIKE 49 |
| 22 | AT5G07030 | AT5G07030 | Eukaryotic aspartyl protease family protein |
| 23 | AT3G54400 | AT3G54400 | Eukaryotic aspartyl protease family protein |
| 24 | WLIM2a | AT2G39900 | GATA type zinc finger transcription factor family protein |
| 25 | AT1G48210 | AT1G48210 | Protein kinase superfamily protein |
| 26 | OEP16-1 | AT2G28900 | outer plastid envelope protein 16-1 |
| 27 | ELF3 | AT2G25930 | hydroxyproline-rich glycoprotein family protein |
| 28 | AT1G54410 | AT1G54410 | dehydrin family protein |
| 29 | TRN2 | AT5G46700 | Tetraspanin family protein |
| 30 | CYS1 | AT5G12140 | cystatin-1 |
| 31 | ADS2 | AT2G31360 | 16:0delta9 desaturase 2 |
| 32 | PWD | AT5G26570 | chloroplastidic phosphoglucan, water dikinase (ATGWD3) |
| 33 | scpl35 | AT5G08260 | serine carboxypeptidase-like 35 |
| 34 | AT4G20430 | AT4G20430 | Subtilase family protein |
| 35 | AT3G58570 | AT3G58570 | P-loop containing nucleoside triphosphate hydrolases superfamily protein |
| 36 | AAP3 | AT1G77380 | amino acid permease 3 |
| 37 | AT5G23575 | AT5G23575 | Transmembrane CLPTM1 family protein |
| 38 | AT1G75060 | AT1G75060 | histone deacetylase complex subunit |
| 39 | AT2G28790 | AT2G28790 | Pathogenesis-related thaumatin superfamily protein |
| 40 | AT1G26450 | AT1G26450 | Carbohydrate-binding X8 domain superfamily protein |
| 41 | VAMP713 | AT5G11150 | vesicle-associated membrane protein 713 |
| 42 | AT2G33830 | AT2G33830 | Dormancy/auxin associated family protein |
| 43 | AT2G21130 | AT2G21130 | Cyclophilin-like peptidyl-prolyl cis-trans isomerase family protein |
| 44 | AT5G48480 | AT5G48480 | Lactoylglutathione lyase / glyoxalase I family protein |
| 45 | TMT2 | AT4G35300 | tonoplast monosaccharide transporter2 |

| Rank | Gene | AGI locus | Annotation |
|---|---|---|---|
| 46 | AT1G54217 | AT1G54217 | Ribosomal protein L18ae family |
| 47 | XTH28 | AT1G14720 | xyloglucan endotransglucosylase/hydrolase 28 |
| 48 | MT2A | AT3G09390 | metallothionein 2A |
| 49 | DYL1 | AT1G28330 | dormancy-associated protein-like 1 |
| 50 | AT5G62200 | AT5G62200 | Embryo-specific protein 3, (ATS3) |

* The 50 highest-ranked co-expression genes retrieved from ATTED-II (v11) for each gene, in the order returned by the database for the Arabidopsis thaliana ath-u.5 dataset. These are the gene sets used for the Gene Ontology enrichment analysis in Table S5 and Fig. S3A, and the networks drawn in Fig. S2. Rank is ATTED-II's ordering; gene symbols and annotations are as supplied by ATTED-II.

**Table S5 |** g:Profiler functional enrichment of the ATTED-II co-expression partners.

### *AtPDF1.2a*

| Source | Term name | Term ID | Adjusted P | Term size | Genes in set | In Fig. S3A | Intersecting genes |
|---|---|---|---|---|---|---|---|
| GO:MF | heme binding | GO:0020037 | 5.21e-03 | 382 | 6 | | AT3G26200, AT5G58390, AT3G26320, AT4G11290, AT3G49110, AT5G36220 |
| GO:MF | catalytic activity | GO:0003824 | 6.88e-03 | 9237 | 28 | | AT5G61160, AT3G04720, AT3G49620, AT2G39030, AT4G16260, AT1G15125, AT3G26200, AT4G24350, AT4G27300, AT2G43590, AT1G07260, AT3G28930, AT5G58390, AT5G11610, AT1G19670, AT3G26320, AT3G57240, AT5G37850, AT4G11290, AT2G02930, AT3G49110, AT5G03630, AT5G36220, AT3G55970, AT1G52400, AT3G14310, AT2G26560, AT1G11350 |
| GO:MF | tetrapyrrole binding | GO:0046906 | 8.61e-03 | 418 | 6 | | AT3G26200, AT5G58390, AT3G26320, AT4G11290, AT3G49110, AT5G36220 |
| GO:MF | molecular_function | GO:0003674 | 3.04e-02 | 22878 | 46 | | AT2G26020, AT5G44430, AT2G26010, AT5G61160, AT3G04720, AT3G23550, AT1G06160, AT3G49620, AT2G39030, AT4G16260, AT1G15125, AT3G26200, AT4G24350, AT4G27300, AT3G16530, AT2G43590, AT3G15356, AT1G07260, AT3G28930, AT5G58390, AT2G32400, AT5G11610, AT2G31230, AT1G19670, AT3G26320, AT2G15080, AT3G57240, AT2G15042, AT5G37850, AT4G11290, AT2G42610, AT2G02930, AT2G14610, AT2G39330, AT3G49110, AT5G03630, AT1G75040, AT5G36220, AT3G55970, AT1G66100, AT4G06746, AT1G52400, AT3G14310, AT5G41761, AT2G26560, AT1G11350 |
| GO:MF | lactoperoxidase activity | GO:0140825 | 4.58e-02 | 76 | 3 | | AT5G58390, AT4G11290, AT3G49110 |
| GO:BP | biological process involved in interspecies interaction between organisms | GO:0044419 | 8.42e-17 | 1477 | 24 | | AT2G26020, AT5G44430, AT2G26010, AT3G04720, AT3G23550, AT1G06160, AT4G16260, AT3G16530, AT3G15356, AT1G07260, AT3G28930, AT1G19670, AT3G57240, AT2G02930, AT2G14610, AT3G49110, AT1G75040, AT3G55970, AT1G66100, AT1G52400, AT3G14310, AT2G26560, AT4G12490, AT1G11350 |
| GO:BP | response to external biotic stimulus | GO:0043207 | 1.28e-15 | 1461 | 23 | | AT2G26020, AT5G44430, AT2G26010, AT3G04720, AT3G23550, AT1G06160, AT4G16260, AT3G16530, AT3G15356, AT1G07260, AT3G28930, AT1G19670, AT3G57240, AT2G02930, AT2G14610, AT3G49110, AT1G75040, AT3G55970, AT1G52400, AT3G14310, AT2G26560, AT4G12490, AT1G11350 |
| GO:BP | response to other organism | GO:0051707 | 1.28e-15 | 1461 | 23 | Yes | AT2G26020, AT5G44430, AT2G26010, AT3G04720, AT3G23550, AT1G06160, AT4G16260, AT3G16530, AT3G15356, AT1G07260, AT3G28930, AT1G19670, AT3G57240, AT2G02930, AT2G14610, AT3G49110, AT1G75040, AT3G55970, AT1G52400, AT3G14310, AT2G26560, AT4G12490, AT1G11350 |
| GO:BP | response to biotic stimulus | GO:0009607 | 1.91e-15 | 1488 | 23 | | AT2G26020, AT5G44430, AT2G26010, AT3G04720, AT3G23550, AT1G06160, AT4G16260, AT3G16530, AT3G15356, AT1G07260, AT3G28930, AT1G19670, AT3G57240, AT2G02930, AT2G14610, AT3G49110, AT1G75040, AT3G55970, AT1G52400, AT3G14310, AT2G26560, AT4G12490, AT1G11350 |
| GO:BP | defense response | GO:0006952 | 1.52e-14 | 1636 | 23 | Yes | AT2G26020, AT5G44430, AT2G26010, AT3G04720, AT3G23550, AT1G06160, AT2G39030, AT4G16260, AT2G43590, AT3G15356, AT1G07260, AT2G31230, AT1G19670, AT2G14610, AT3G49110, AT1G75040, AT3G55970, AT1G66100, AT1G52400, AT3G14310, AT2G26560, AT4G12490, AT1G11350 |

| Source | Term name | Term ID | Adjusted P | Term size | Genes in set | In Fig. S3A | Intersecting genes |
|---|---|---|---|---|---|---|---|
| GO:BP | response to external stimulus | GO:0009605 | 2.85e-14 | 1684 | 23 | | AT2G26020, AT5G44430, AT2G26010, AT3G04720, AT3G23550, AT1G06160, AT4G16260, AT3G16530, AT3G15356, AT1G07260, AT3G28930, AT1G19670, AT3G57240, AT2G02930, AT2G14610, AT3G49110, AT1G75040, AT3G55970, AT1G52400, AT3G14310, AT2G26560, AT4G12490, AT1G11350 |
| GO:BP | response to stimulus | GO:0050896 | 9.02e-13 | 6002 | 35 | | AT2G26020, AT5G44430, AT2G26010, AT3G04720, AT3G23550, AT1G06160, AT2G39030, AT4G16260, AT4G24350, AT3G16530, AT2G43590, AT3G15356, AT1G07260, AT3G28930, AT5G58390, AT2G32400, AT2G31230, AT1G19670, AT3G57240, AT5G37850, AT3G16670, AT4G11290, AT2G02930, AT2G14610, AT3G49110, AT5G03630, AT1G75040, AT3G55970, AT1G66100, AT4G06746, AT1G52400, AT3G14310, AT2G26560, AT4G12490, AT1G11350 |
| GO:BP | response to stress | GO:0006950 | 1.27e-11 | 3733 | 28 | | AT2G26020, AT5G44430, AT2G26010, AT3G04720, AT3G23550, AT1G06160, AT2G39030, AT4G16260, AT4G24350, AT2G43590, AT3G15356, AT1G07260, AT5G58390, AT2G31230, AT1G19670, AT5G37850, AT3G16670, AT4G11290, AT2G14610, AT3G49110, AT1G75040, AT3G55970, AT1G66100, AT1G52400, AT3G14310, AT2G26560, AT4G12490, AT1G11350 |
| GO:BP | defense response to fungus | GO:0050832 | 2.73e-08 | 576 | 12 | Yes | AT2G26020, AT5G44430, AT2G26010, AT3G04720, AT4G16260, AT3G15356, AT1G19670, AT3G49110, AT3G55970, AT1G52400, AT3G14310, AT4G12490 |
| GO:BP | response to fungus | GO:0009620 | 1.39e-07 | 664 | 12 | Yes | AT2G26020, AT5G44430, AT2G26010, AT3G04720, AT4G16260, AT3G15356, AT1G19670, AT3G49110, AT3G55970, AT1G52400, AT3G14310, AT4G12490 |
| GO:BP | response to chemical | GO:0042221 | 2.60e-07 | 2941 | 21 | | AT3G04720, AT3G23550, AT1G06160, AT2G39030, AT3G16530, AT3G15356, AT5G58390, AT2G32400, AT2G31230, AT4G11290, AT2G02930, AT2G14610, AT3G49110, AT5G03630, AT1G75040, AT3G55970, AT4G06746, AT1G52400, AT3G14310, AT2G26560, AT1G11350 |
| GO:BP | defense response to other organism | GO:0098542 | 6.57e-05 | 387 | 8 | | AT3G04720, AT3G23550, AT1G06160, AT1G07260, AT2G14610, AT3G49110, AT1G75040, AT2G26560 |
| GO:BP | defense response to symbiont | GO:0140546 | 3.29e-04 | 325 | 7 | | AT3G04720, AT1G06160, AT1G07260, AT2G14610, AT3G49110, AT1G75040, AT2G26560 |
| GO:BP | cellular response to chemical stimulus | GO:0070887 | 3.13e-03 | 1670 | 12 | | AT1G06160, AT3G15356, AT5G58390, AT2G32400, AT2G31230, AT4G11290, AT3G49110, AT3G55970, AT4G06746, AT1G52400, AT2G26560, AT1G11350 |
| GO:BP | response to bacterium | GO:0009617 | 3.41e-03 | 463 | 7 | | AT3G04720, AT3G28930, AT1G19670, AT3G57240, AT3G49110, AT3G14310, AT1G11350 |
| GO:BP | systemic acquired resistance | GO:0009627 | 6.00e-03 | 90 | 4 | Yes | AT3G04720, AT1G07260, AT2G14610, AT1G75040 |
| GO:BP | detoxification | GO:0098754 | 3.42e-02 | 276 | 5 | | AT3G23550, AT5G58390, AT4G11290, AT2G02930, AT3G49110 |
| GO:BP | response to ethylene | GO:0009723 | 3.79e-02 | 282 | 5 | Yes | AT3G04720, AT1G06160, AT3G15356, AT2G31230, AT4G06746 |
| GO:CC | secretory vesicle | GO:0099503 | 4.00e-10 | 170 | 9 | | AT2G26010, AT3G04720, AT4G16260, AT2G43590, AT3G57240, AT4G11290, AT2G14610, AT3G49110, AT1G75040 |
| GO:CC | extracellular region | GO:0005576 | 3.68e-07 | 1772 | 16 | | AT2G26020, AT5G44430, AT2G26010, AT4G16260, AT3G16530, AT3G15356, AT5G58390, AT3G57240, AT4G11290, AT2G14610, AT3G49110, AT1G75040, AT1G66100, AT3G14310, AT4G12490, AT1G11350 |
| GO:CC | cytoplasmic vesicle | GO:0031410 | 1.47e-04 | 745 | 9 | | AT2G26010, AT3G04720, AT4G16260, AT2G43590, AT3G57240, AT4G11290, AT2G14610, |

| Source | Term name | Term ID | Adjusted P | Term size | Genes in set | In Fig. S3A | Intersecting genes |
|---|---|---|---|---|---|---|---|
| | | | | | | | AT3G49110, AT1G75040 |
| GO:CC | intracellular vesicle | GO:0097708 | 1.50e-04 | 747 | 9 | | AT2G26010, AT3G04720, AT4G16260, AT2G43590, AT3G57240, AT4G11290, AT2G14610, AT3G49110, AT1G75040 |
| GO:CC | vesicle | GO:0031982 | 1.95e-04 | 771 | 9 | | AT2G26010, AT3G04720, AT4G16260, AT2G43590, AT3G57240, AT4G11290, AT2G14610, AT3G49110, AT1G75040 |
| GO:CC | vacuole | GO:0005773 | 3.18e-03 | 1094 | 9 | | AT3G04720, AT4G16260, AT5G58390, AT1G19670, AT4G11290, AT2G39330, AT3G49110, AT1G75040, AT1G52400 |
| GO:CC | apoplast | GO:0048046 | 8.56e-03 | 489 | 6 | | AT3G16530, AT3G15356, AT3G57240, AT2G14610, AT1G75040, AT3G14310 |
| GO:CC | cellular anatomical structure | GO:0110165 | 3.49e-02 | 18497 | 40 | | AT2G26020, AT5G44430, AT2G26010, AT3G04720, AT3G23550, AT1G06160, AT4G16260, AT3G26200, AT4G24350, AT4G27300, AT3G16530, AT2G43590, AT3G15356, AT3G28930, AT5G58390, AT2G32400, AT5G11610, AT2G31230, AT1G19670, AT3G26320, AT2G15080, AT3G57240, AT5G37850, AT4G11290, AT2G42610, AT2G02930, AT2G14610, AT2G39330, AT3G49110, AT5G03630, AT1G75040, AT5G36220, AT1G66100, AT4G06746, AT1G52400, AT3G14310, AT5G41761, AT2G26560, AT4G12490, AT1G11350 |
| KEGG | MAPK signaling pathway - plant | KEGG:04016 | 1.86e-03 | 146 | 4 | | AT2G26020, AT5G44430, AT2G26010, AT2G14610 |
| KEGG | Phenylpropanoid biosynthesis | KEGG:00940 | 2.62e-02 | 130 | 3 | | AT5G58390, AT4G11290, AT3G49110 |

### *AtPDF2.2*

| Source | Term name | Term ID | Adjusted P | Term size | Genes in set | In Fig. S3A | Intersecting genes |
|---|---|---|---|---|---|---|---|
| GO:BP | response to temperature stimulus | GO:0009266 | 2.70e-04 | 644 | 9 | Yes | AT4G38580, AT5G08000, AT3G05880, AT2G21660, AT1G13930, AT2G28900, AT2G25930, AT5G12140, AT4G35300 |
| GO:BP | circadian rhythm | GO:0007623 | 1.39e-03 | 142 | 5 | Yes | AT2G21660, AT5G61380, AT2G40080, AT2G25930, AT1G75060 |
| GO:BP | rhythmic process | GO:0048511 | 1.88e-03 | 151 | 5 | Yes | AT2G21660, AT5G61380, AT2G40080, AT2G25930, AT1G75060 |
| GO:BP | response to stimulus | GO:0050896 | 3.48e-03 | 6002 | 23 | | AT2G02120, AT2G02130, AT4G38580, AT4G30650, AT5G57110, AT5G08000, AT3G05880, AT2G21660, AT5G61380, AT1G13930, AT5G58600, AT2G40080, AT2G28900, AT2G25930, AT5G46700, AT5G12140, AT2G31360, AT1G77380, AT2G28790, AT5G11150, AT2G33830, AT4G35300, AT1G28330 |
| GO:BP | circumnutation | GO:0010031 | 1.85e-02 | 6 | 2 | | AT5G61380, AT2G25930 |
| GO:BP | multicellular organismal movement | GO:0050879 | 1.85e-02 | 6 | 2 | | AT5G61380, AT2G25930 |

| Source | Term name | Term ID | Adjusted P | Term size | Genes in set | In Fig. S3A | Intersecting genes |
|---|---|---|---|---|---|---|---|
| GO:BP | response to cold | GO:0009409 | 1.98e-02 | 410 | 6 | Yes | AT3G05880, AT2G21660, AT2G28900, AT2G25930, AT5G12140, AT4G35300 |
| GO:BP | response to abiotic stimulus | GO:0009628 | 3.53e-02 | 2131 | 12 | Yes | AT4G38580, AT5G08000, AT3G05880, AT2G21660, AT1G13930, AT2G40080, AT2G28900, AT2G25930, AT5G12140, AT2G31360, AT5G11150, AT4G35300 |
| GO:BP | regulation of circadian rhythm | GO:0042752 | 4.24e-02 | 54 | 3 | | AT2G40080, AT2G25930, AT1G75060 |
| GO:BP | response to stress | GO:0006950 | 4.52e-02 | 3733 | 16 | Yes | AT2G02120, AT2G02130, AT4G38580, AT4G30650, AT5G08000, AT3G05880, AT2G21660, AT1G13930, AT5G58600, AT2G28900, AT2G25930, AT5G12140, AT1G77380, AT2G28790, AT5G11150, AT4G35300 |
| GO:CC | cellular anatomical structure | GO:0110165 | 2.80e-06 | 18497 | 46 | | AT2G02120, AT2G02130, AT4G38580, AT4G30650, AT5G57110, AT1G68220, AT5G39410, AT1G08980, AT5G08000, AT2G32280, AT3G05880, AT2G21660, AT5G61380, AT1G13930, AT5G58600, AT1G25320, AT1G53035, AT3G54820, AT4G26670, AT2G40080, AT3G10410, AT3G54400, AT2G39900, AT1G48210, AT2G28900, AT2G25930, AT1G54410, AT5G46700, AT5G12140, AT2G31360, AT5G26570, AT5G08260, AT4G20430, AT1G77380, AT5G23575, AT1G75060, AT2G28790, AT1G26450, AT5G11150, AT2G33830, AT2G21130, AT5G48480, AT4G35300, AT1G54217, AT1G14720, AT5G62200 |
| GO:CC | cellular_component | GO:0005575 | 7.13e-05 | 20029 | 46 | | AT2G02120, AT2G02130, AT4G38580, AT4G30650, AT5G57110, AT1G68220, AT5G39410, AT1G08980, AT5G08000, AT2G32280, AT3G05880, AT2G21660, AT5G61380, AT1G13930, AT5G58600, AT1G25320, AT1G53035, AT3G54820, AT4G26670, AT2G40080, AT3G10410, AT3G54400, AT2G39900, AT1G48210, AT2G28900, AT2G25930, AT1G54410, AT5G46700, AT5G12140, AT2G31360, AT5G26570, AT5G08260, AT4G20430, AT1G77380, AT5G23575, AT1G75060, AT2G28790, AT1G26450, AT5G11150, AT2G33830, AT2G21130, AT5G48480, AT4G35300, AT1G54217, AT1G14720, AT5G62200 |
| GO:CC | cell periphery | GO:0071944 | 3.36e-04 | 2917 | 16 | | AT4G38580, AT5G57110, AT5G39410, AT5G08000, AT1G13930, AT1G25320, AT3G54820, AT1G48210, AT5G12140, AT1G77380, AT2G28790, AT1G26450, AT5G11150, AT2G21130, AT4G35300, AT1G14720 |
| GO:CC | plasma membrane | GO:0005886 | 1.43e-03 | 2524 | 14 | | AT4G38580, AT5G57110, AT5G39410, AT5G08000, AT1G13930, AT1G25320, AT3G54820, AT1G48210, AT5G12140, AT1G77380, AT1G26450, AT5G11150, AT2G21130, AT4G35300 |
| GO:CC | membrane | GO:0016020 | 3.04e-03 | 8169 | 26 | | AT4G38580, AT4G30650, AT5G57110, AT1G68220, AT5G39410, AT5G08000, AT2G32280, AT3G05880, AT1G13930, AT5G58600, AT1G25320, AT1G53035, AT3G54820, AT4G26670, AT1G48210, AT2G28900, AT5G46700, AT5G12140, AT2G31360, AT1G77380, AT5G23575, AT1G26450, AT5G11150, AT2G21130, AT4G35300, AT1G54217 |
| GO:CC | extracellular region | GO:0005576 | 5.16e-03 | 1772 | 11 | | AT2G02120, AT2G02130, AT1G08980, AT2G21660, AT1G13930, AT3G10410, AT3G54400, AT5G08260, AT4G20430, AT1G14720, AT5G62200 |
| GO:CC | cell junction | GO:0030054 | 5.30e-03 | 900 | 8 | | AT2G02130, AT4G38580, AT5G57110, AT5G08000, AT2G21660, AT5G46700, AT2G28790, AT1G26450 |
| GO:CC | cell-cell junction | GO:0005911 | 5.30e-03 | 900 | 8 | | AT2G02130, AT4G38580, AT5G57110, AT5G08000, AT2G21660, AT5G46700, AT2G28790, AT1G26450 |
| GO:CC | anchoring junction | GO:0070161 | 5.30e-03 | 900 | 8 | | AT2G02130, AT4G38580, AT5G57110, AT5G08000, AT2G21660, AT5G46700, AT2G28790, AT1G26450 |

| Source | Term name | Term ID | Adjusted P | Term size | Genes in set | In Fig. S3A | Intersecting genes |
|---|---|---|---|---|---|---|---|
| GO:CC | plasmodesma | GO:0009506 | 5.30e-03 | 900 | 8 | | AT2G02130, AT4G38580, AT5G57110, AT5G08000, AT2G21660, AT5G46700, AT2G28790, AT1G26450 |
| GO:CC | symplast | GO:0055044 | 5.35e-03 | 901 | 8 | | AT2G02130, AT4G38580, AT5G57110, AT5G08000, AT2G21660, AT5G46700, AT2G28790, AT1G26450 |
| GO:CC | intracellular anatomical structure | GO:0005622 | 9.89e-03 | 13505 | 34 | | AT2G02130, AT4G38580, AT5G57110, AT5G39410, AT1G08980, AT2G32280, AT2G21660, AT5G61380, AT1G13930, AT5G58600, AT4G26670, AT2G40080, AT3G10410, AT3G54400, AT2G39900, AT1G48210, AT2G28900, AT2G25930, AT1G54410, AT5G12140, AT2G31360, AT5G26570, AT1G77380, AT5G23575, AT1G75060, AT2G28790, AT1G26450, AT5G11150, AT2G33830, AT2G21130, AT5G48480, AT4G35300, AT1G54217, AT5G62200 |
| GO:CC | intracellular organelle | GO:0043229 | 1.14e-02 | 12312 | 32 | | AT2G02130, AT4G38580, AT5G57110, AT5G39410, AT1G08980, AT2G32280, AT2G21660, AT5G61380, AT1G13930, AT5G58600, AT4G26670, AT2G40080, AT3G10410, AT2G39900, AT1G48210, AT2G28900, AT2G25930, AT1G54410, AT5G12140, AT2G31360, AT5G26570, AT1G77380, AT5G23575, AT1G75060, AT2G28790, AT1G26450, AT5G11150, AT2G33830, AT2G21130, AT4G35300, AT1G54217, AT5G62200 |
| GO:CC | organelle | GO:0043226 | 1.17e-02 | 12326 | 32 | | AT2G02130, AT4G38580, AT5G57110, AT5G39410, AT1G08980, AT2G32280, AT2G21660, AT5G61380, AT1G13930, AT5G58600, AT4G26670, AT2G40080, AT3G10410, AT2G39900, AT1G48210, AT2G28900, AT2G25930, AT1G54410, AT5G12140, AT2G31360, AT5G26570, AT1G77380, AT5G23575, AT1G75060, AT2G28790, AT1G26450, AT5G11150, AT2G33830, AT2G21130, AT4G35300, AT1G54217, AT5G62200 |
| GO:CC | vacuole | GO:0005773 | 2.03e-02 | 1094 | 8 | | AT5G39410, AT2G21660, AT3G10410, AT2G28900, AT5G12140, AT5G11150, AT4G35300, AT5G62200 |
| GO:CC | intracellular membrane-bounded organelle | GO:0043231 | 4.05e-02 | 11808 | 30 | | AT2G02130, AT4G38580, AT5G57110, AT5G39410, AT1G08980, AT2G32280, AT2G21660, AT5G61380, AT1G13930, AT5G58600, AT4G26670, AT2G40080, AT3G10410, AT1G48210, AT2G28900, AT2G25930, AT1G54410, AT5G12140, AT2G31360, AT5G26570, AT1G77380, AT5G23575, AT1G75060, AT2G28790, AT1G26450, AT5G11150, AT2G33830, AT2G21130, AT4G35300, AT5G62200 |
| GO:CC | membrane-bounded organelle | GO:0043227 | 4.15e-02 | 11823 | 30 | | AT2G02130, AT4G38580, AT5G57110, AT5G39410, AT1G08980, AT2G32280, AT2G21660, AT5G61380, AT1G13930, AT5G58600, AT4G26670, AT2G40080, AT3G10410, AT1G48210, AT2G28900, AT2G25930, AT1G54410, AT5G12140, AT2G31360, AT5G26570, AT1G77380, AT5G23575, AT1G75060, AT2G28790, AT1G26450, AT5G11150, AT2G33830, AT2G21130, AT4G35300, AT5G62200 |
| KEGG | Circadian rhythm - plant | KEGG:04712 | 4.98e-02 | 43 | 2 | | AT5G61380, AT2G25930 |

* Gene Ontology and KEGG enrichment of the 50-gene co-expression sets in Table S4, computed with g:Profiler using the g:SCS multiple-testing correction at a threshold of 0.05 against the Arabidopsis thaliana annotation (query size 50; effective domain size 32,833). All significant terms are listed. Terms plotted in Fig. S3A are marked in the "In Fig. S3A" column.
